# Membrane lipid poly-unsaturation selectively affects dopamine D2 receptor internalization

**DOI:** 10.1101/2023.12.14.571632

**Authors:** Silvia Sposini, Rim Baccouch, Mathias Lescuyer, Aditi A Mali, Véronique De Smedt-Peyrusse, Joyce Heuninck, Ana Gorse, Thierry Durroux, Pierre Trifilieff, David Perrais, Isabel Alves

## Abstract

The brain is highly enriched in poly-unsaturated fatty acids (PUFAs) and their deficiency has been associated with several neuropsychiatric disorders. Here, we demonstrate that the Dopamine receptor D2 (D2R), a class A G protein coupled receptor (GPCR) which is a main target of antipsychotics, displays unique sensitivity to membrane PUFA composition. We found that membrane enrichment with either of two distinct PUFAs significantly impairs agonist-induced D2R endocytosis in HEK-293 cells and cortical neurons. This treatment does not affect clathrin-mediated endocytosis or the internalization of several other GPCRs. Moreover, we show that D2R clustering at endocytic pits is not affected, but that recruitment of β-arrestin2 is strongly impaired and endocytic vesicle formation is slowed down. Finally, mutation of key residues in intracellular loop 2 abolishes the sensitivity of D2R endocytosis to PUFA enrichment. We conclude that D2R trafficking is uniquely dependent on membrane PUFAs, which could influence its role in the control of brain function and behavior.

## Introduction

Cellular signaling events typically involve complex interactions between protein dynamics and membrane remodeling processes. In particular, lipids are key regulators of membrane protein structure, conformational flexibility and activity. For example, in reconstituted *in vitro* systems, the activity of many G protein coupled receptors (GPCRs), the largest family of membrane receptors, such as rhodopsin, the serotonin 5HT_1A_ receptor or the chemokine CCR5 receptor, is strongly influenced by the presence of cholesterol in the membrane ^1–4^. Moreover, the nature of phospholipid head groups (phosphatidylcholine or phosphatidylethanolamine) also affects the binding of agonists or antagonists to GPCRs and their activation such as for the β2 adrenergic receptor (β2AR) ^5^. Finally, the nature of the phospholipid fatty acid chains may also affect GPCR activity. Indeed, poly-unsaturated fatty acids (PUFAs) are necessary for preserving the optimal conformation and activity of GPCRs like rhodopsin and adenosine A_2A_ receptors ^6–9^.

PUFAs represent a class of fatty acids characterized by two or more double bonds within their carbon chains. The position of the first carbon possessing an unsaturation, counted from the methyl end, known as omega, defines the type of PUFA, e.g. ω3 for first unsaturation at the third carbon atom from the methyl end. During brain development, PUFAs accumulate perinatally and ultimately comprise approximately 30 % of total brain fatty acids in adulthood. They modulate membrane properties, signaling pathways, and cell metabolism ^10^. Deficits in brain PUFA accumulation have been linked to several psychiatric disorders including major depressive disorder, bipolar disorder or schizophrenia ^11,12^. In particular, the brain of individuals suffering from psychiatric disorders often display lower levels of ω3-PUFAs ^13–15^ and some studies have reported a significant advantage of ω3-PUFAs dietary supplementation for reducing symptoms ^16,17^. However, the molecular mechanisms by which PUFAs participate in the etiology of psychiatric symptoms remain largely elusive.

A shared neurobiological feature observed across multiple psychiatric disorders involves dysregulation of dopamine neurotransmission, in particular, the activity of the dopamine D2 receptor (D2R) ^18,19^. Interestingly, ω3-PUFA deficiency in animal models have revealed alterations of dopamine transmission ^20–22^ and a particular vulnerability of D2R-expressing neurons to ω3-PUFAs deficiency ^23^. Mechanistically, molecular dynamics simulations show that membrane PUFA levels modulate the membrane diffusion rate of the D2R ^24^. Moreover, enrichment in PUFAs in HEK-293 cells alters D2R ligand binding and signaling ^25^. Altogether, these findings suggest a direct link between membrane PUFAs and the activity of the D2R.

Because the signaling efficiency mediated by D2R is directly related to the overall number of responsive receptors at the cellular membrane, which is controlled by desensitization and internalization processes, membrane PUFA composition, by affecting membrane biophysical properties ^26,27^ or GPCR conformation, could affect its trafficking. Activation by agonists leads to D2R phosphorylation by G protein receptor kinase 2/3 (GRK 2/3) on serine and threonine residues in the second and third intracellular loops (ICL2/3)^28^. This phosphorylation triggers robust recruitment of the cytoplasmic protein ꞵ-arrestin2, which binds and targets the phosphorylated receptor by scaffolding it to clathrin-coated pits (CCPs), leading to its internalization into endosomes after dynamin-dependent membrane scission ^29–32^. Furthermore, ꞵ-arrestins also function as scaffold proteins that interact with cytoplasmic proteins and link GPCRs to intracellular signaling pathways such as MAPK cascades ^33–35^.

Here we investigated the modulatory effect of membrane lipid poly-unsaturation on ligand-induced D2R internalization. We treated D2R-expressing HEK-293 cells or cortical neurons in culture with either the ω3-PUFA docosahexaenoic acid (DHA) or the ω6-PUFA docosapentaenoic acid (DPA) in order to increase membrane poly-unsaturation in a controlled manner. Using resonance energy transfer-based technique or confocal microscopy, we demonstrated that PUFA membrane enrichment strongly blunts ligand-induced internalization of D2R but does not affect the internalization of transferrin, a cargo of clathrin mediated endocytosis (CME), or 7 other class A GPCRs. In addition, we found using Total Internal Reflection fluorescence (TIRF) microscopy that ligand-induced clustering of D2R is preserved in PUFA-enriched cells but that β-arrestin2 membrane recruitment is decreased and endocytic vesicle formation is reduced in conditions of PUFA enrichment. Finally, we show that the unique sensitivity of D2R to PUFAs depends on two membrane proximal serine residues located in ICL2 of D2R.

## Results

### Robust incorporation of exogenous fatty acids into cell membranes

To assess the potential effects of membrane PUFAs on D2R trafficking, we used the protocol developed in ^25^ to enrich membrane phospholipids with selected fatty acids (FAs). We evaluated the consequences of enrichment with 2 PUFAs, namely DHA or DPA, and the saturated fatty acid behenic acid (BA), all displaying 22 carbons (Figure 1A). This allows the distinction between the effect of number and position of unsaturations *versus* carbon chain length. We first tested the maximum dose of exogenous FA tolerated by cells. Incubation for 24 h with 30 µM of any FA does not affect cell viability. However, incubation of DHA at 60 µM significantly reduces cell viability (Figure 1B). Therefore, we used 30 µM FA treatment for 24 h to get maximal effects. At these doses, FA treatment does not affect the expression of D2R at the cell surface (Figure 1C). Incubation with either DHA or DPA significantly increases PUFA levels in phospholipids, from 8.6 % in control (EtOH) to ∼14 %, while incubation with 30 µM of BA does not affect it significantly (Figure 1D). Conversely, the content of saturated FAs in phospholipids does not change after DHA or DPA treatment, while it increases in BA treated cells (Figure 1E). Incubation with either FA (DHA, DPA or BA) increases its respective content in phospholipids by at least 3 fold, leaving the content of the other FAs unaffected (Figure 1F-H). Consequently, the ratio of total ω6 (such as DPA) and ω3 (such as DHA) PUFAs (ω6/ω3) is significantly changed for cells treated with DHA (3-fold lower, p < 0.01) or DPA (2.7-fold higher, p < 0.0001), but not for cells treated with BA (Figure 1H). This shows that these treatments alter the cellular phospholipid composition in a FA specific and controlled manner.

**Figure 1:**
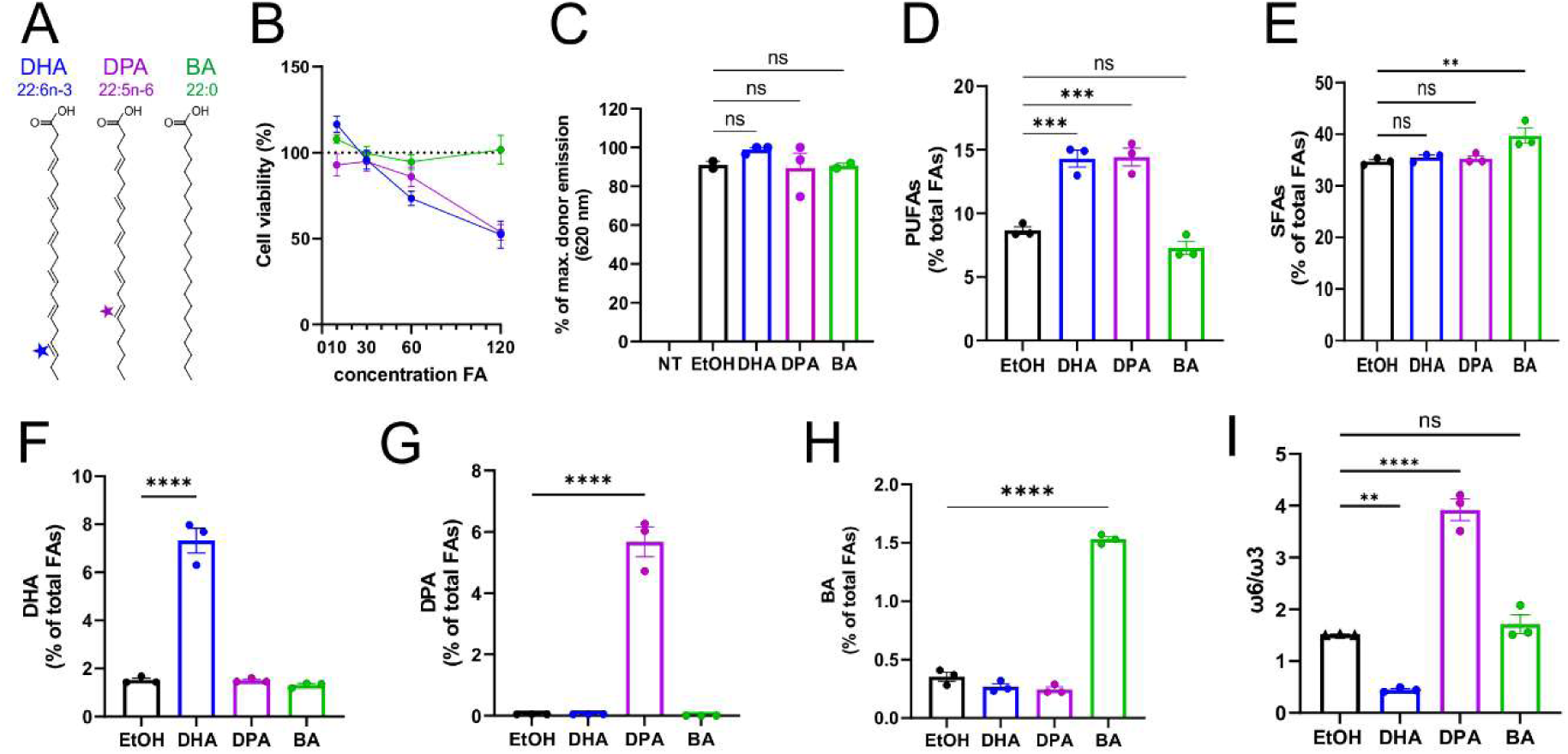
Phospholipid fatty acid enrichment in cells treated with exogenous fatty acids. **A:** Fatty acid structures: DHA (Docosahexaenoic acid), an ω3-PUFA with the formula C_22_H_32_O_2_; DPA (Docosapentaenoic acid), an ω6-PUFA with the formula C_22_H_34_O_2_. BA (behenic acid), a saturated fatty acid (SFA) with the formula C_22_H_44_O_2_. **B:** Viability of cells treated with FAs (BA, DHA or DPA), normalized to cells incubated with vehicle (0.4 % ethanol). **C:** D2R membrane expression levels measured by the SNAP-Lumi-4-Tb fluorescence intensity at 620 nm as for the DERET assays (see Figure 2) in non-transfected (NT) or cells co-expressing SNAP-D2R, β-arrestin2 and GRK2 treated with EtOH or FAs (BA, DHA, DPA). **D-E:** Cellular amount of total PUFAs (B) and total SFAs (C), expressed as percentage of total phospholipid fatty acid (FA), measured in control (EtOH) and FAs (BA, DHA or DPA) treated cells (30 µM, 24 h). Control cells were treated in presence of 0.1 % of ethanol (EtOH, solvent used in BA and PUFAs solubilisation). **F-H:** Cellular amount of DHA (D), DPA (E) and BA (F), expressed as percentage of total phospholipid fatty acid, measured in control (EtOH) and cells treated with the corresponding fatty acid. **I:** Ratio of total ω6 and ω3 PUFAs (ω6/ω3) measured in control (EtOH) and fatty acid treated cells. One-way ANOVA with Dunnett’s multiple comparisons test of 3 independent experiments; ns p ≥ 0.05, * p<0.05, ** p<0.001, *** p<0.005, **** p<0.001.

### Impact of PUFA enrichment on D2R internalization investigated by DERET assay

We assessed the effect of PUFA enrichment on the internalization of D2R induced by multiple ligands in a dose-response manner using a high-throughput fluorometric assay, diffusion-enhanced resonance energy transfer (DERET). This assay measures the transfer of energy between the long life time luminescence donor Lumi-4-terbium (measured at 620 nm) attached to SNAP tagged D2R (SNAP-D2R) and the extracellular fluorescence acceptor fluorescein (peak fluorescence 520 nm) ^36^. As illustrated in Figure S1A, agonist-induced SNAP-D2R internalization results in a loss of energy transfer, leading to an increased fluorescence Ratio (620/520). In HEK-293 cells transfected with SNAP-D2R alone, addition of the D2R agonist quinpirole (QPL, 10 µM) increases the Ratio less than 2-fold in 40 minutes compared to cells without agonist. Co-transfection of cells with GRK2, which phosphorylates D2R, and βarrestin2, which binds phosphorylated D2R, increases receptor internalization 4 to 6 fold. We obtained a maximal internalization in cells co-transfected with GRK2 and βarrestin2 (Figure S1B). Therefore, we used throughout this study HEK-293 cells co-transfected with tagged D2R, GRK2 and βarrestin2. To further confirm that increase in DERET Ratio reflects endocytosis, we applied in succession dopamine (10 µM), which increases ratio, then dopamine plus the D2R antagonist haloperidol (10 µM) or the partial agonist aripiprazole (100 μM), which reversed the increases, showing recycling back to the plasma membrane of labelled receptors (Figure S1C). Taken together, these results show that DERET assay is a suitable technique to monitor agonist-induced D2R internalization in real time.

We next investigated the impact of DHA, DPA or BA enrichment on agonist-induced D2R internalization. Importantly, these treatments do not affect the level of D2R surface expression, as measured by SNAP-Lumi4-Tb fluorescence intensity at 620 nm (donor fluorescence) (Figure 1C). However, enrichment with DHA or DPA strongly blunts D2R internalization induced by either dopamine or QPL (Figure 2A, B). We observed this effect of PUFAs for all tested agonist concentrations. Dose-response curves, taken after 30 min stimulation, reveal a significant reduction of the maximal response (E_max_) for PUFA-enriched cells (Figures 2C-E). However, we observed no significant difference in dopamine-induced internalization efficacy (EC_50_) under membrane PUFA enrichment, while EC_50_ for QPL is significantly higher when compared to control cells (Figure 2F). We obtained similar dose-response curves with longer incubations (45, 60 or 75 min) with either dopamine or QPL (Figure S2). Conversely, treatment with BA does not affect D2R internalization induced by either dopamine or QPL (10 µM) (Figure 1G, H). Altogether, these data indicate that the decrease in D2R internalization observed in membrane PUFA enriched cells is not due to the increased amount of long chain fatty acids (C22) but rather to the increased levels of membrane poly-unsaturation, independently of the position of the first unsaturation.

**Figure 2:**
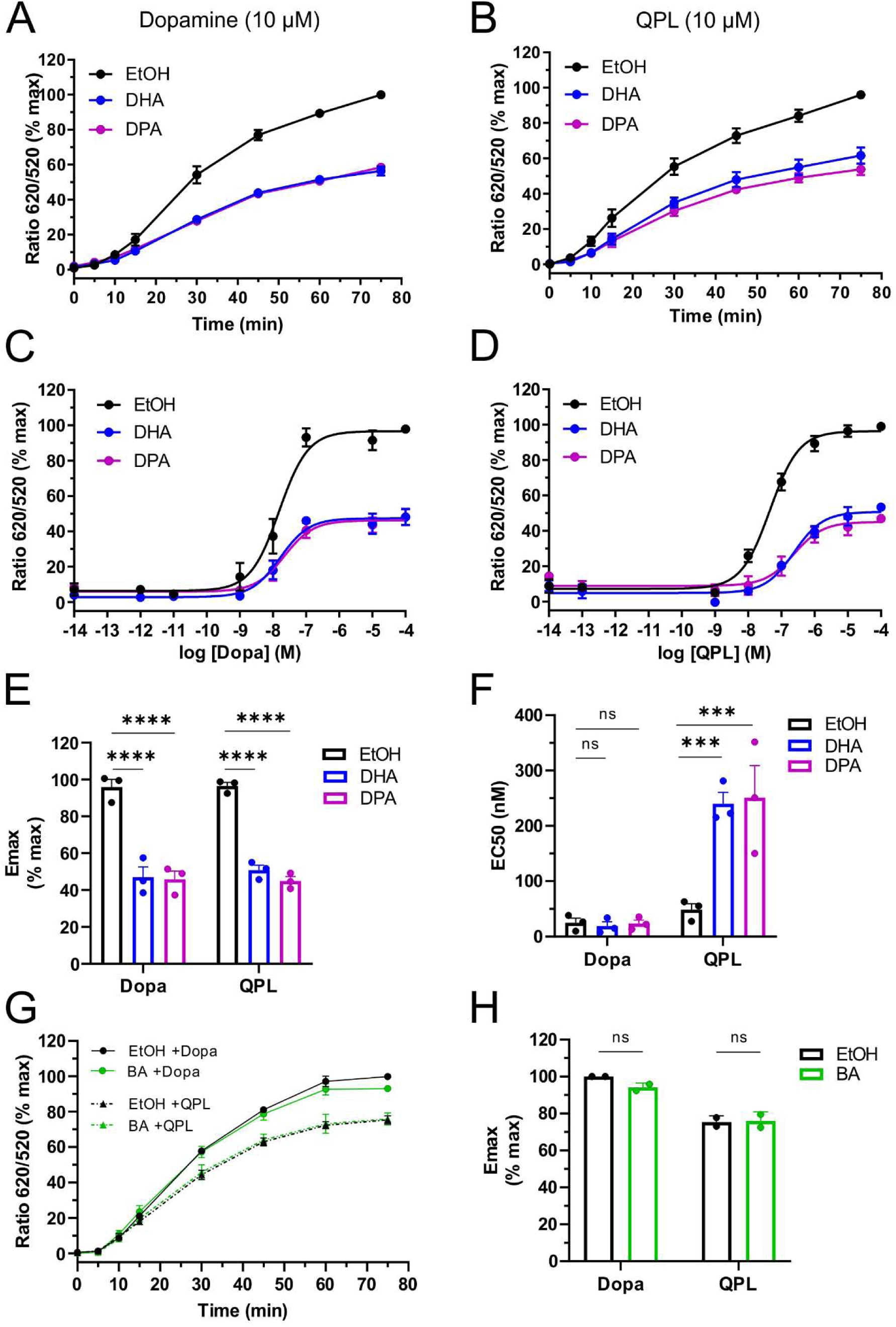
Impact of PUFA enrichment on D2R induced internalization assessed by DERET. **A, B**: Real-time internalization of SNAP tagged D2R upon Dopamine (A) or Quinpirole (B) stimulation (10 µM) of control (EtOH) and PUFA (DHA or DPA) enriched cells. Percentage of fluorescence ratio (ratio of the donor/acceptor fluorescence intensity; 620/520 nm) were plotted as a function of time. **C, D:** Dose–response curves of D2R agonist (Dopa, C; QPL, D) induced internalization following 30 min incubation at 37°C in presence of increasing concentrations of the indicated agonist. Data were fitted using non-linear regression dose–response and expressed as log [ligand] *versus* response for the 3 different conditions. **E, F:** Emax and EC_50_ values obtained from dose-response experiments of D2R internalization (C and D). **G:** Real-time internalization of SNAP tagged D2R expressed in control (EtOH) and BA-enriched cells upon stimulation with 10 µM of dopamine or quinpirole. **H**: Emax values measured in control and BA-enriched cells 30 min after dopamine or quinpirole stimulation. 3 independent experiments carried out in triplicates. Two-way ANOVA test with Sidak’s multiple comparisons test; **** p <0.0001, *** p <0.001, ns p ≥ 0.05.

### PUFA levels selectively affect D2R internalization without affecting general endocytosis

To confirm the effect of DHA and DPA on D2R endocytosis, we examined the formation of endosomes with confocal microscopy. Cells expressing FLAG-D2R, GRK2 and β-arrestin2 were labelled live with anti-FLAG antibody for 10 min, incubated with or without D2R agonists (dopamine or QPL) for 30 min and fixed. As shown in Figures 3A and C, in the absence of D2R agonists, FLAG staining was mostly restricted to the cell perimeter, suggesting that the D2R has little constitutive internalization in all conditions of PUFA treatment. Addition of dopamine for 30 min triggered the formation of punctuate structures corresponding to internalized receptors. In DHA and DPA treated cells, the number of puncta induced by dopamine was much smaller than in untreated cells (34.5 ± 6.12 % and 29.5 ± 6 % for DHA and DPA compared to EtOH, respectively, p < 0.0001) (Figure 3A, B). We obtained similar results in cells treated with QPL (61.8 ± 6.9 % of control for DHA and 48.5 ± 7.3 % DPA, respectively, p < 0.0001) (Figure 3C, D). These results further show that cell enrichment with each PUFA dampens agonist-induced D2R internalization.

**Figure 3:**
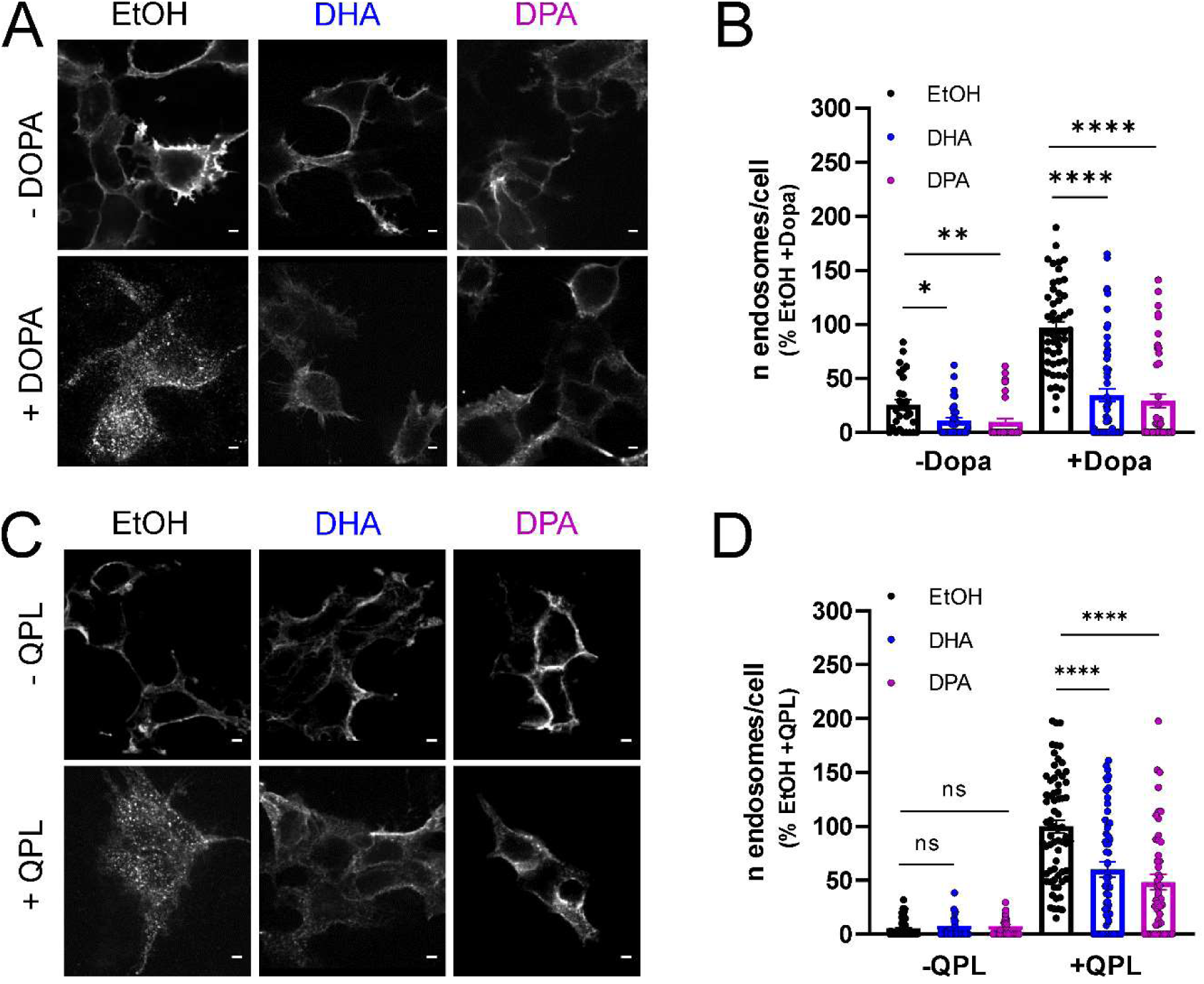
PUFA enrichment affects D2R internalization. **A, C:** Representative confocal microscopy images of control (EtOH) or PUFA-enriched (DHA or DPA) HEK-293 cells expressing FLAG-D2R, GRK2 and βarr2, before or after stimulation with either 10 µM of dopamine (Dopa) (A) or 10 µM of quinpirole (QPL) (C) for 30 min. Scale bar: 5 μm. **B, D:** Quantification of the number of D2R puncta (endosomes) generated before and after dopamine (Dopa) (B) or quinpirole (QPL) (D) stimulation in control (EtOH) and PUFA enriched cells. 68-71 cells per condition collected across 3 independent experiments. One-way ANOVA with Dunnett’s multiple comparisons tests; * *p* < 0.0001, ** *p* < 0.005, **** *p* < 0.0001, ns *p* ≥ 0.05.

D2Rs are internalized through CME ^32^. We thus tested if PUFA treatment affected the internalization of transferrin, a prototypical CME cargo ^37^. Fluorescently-labeled transferrin (Tfn-A568) uptake was assessed by confocal microscopy in both control and PUFA-enriched cells, incubated for 5 min at 37 °C in presence of Tfn-A568. As shown if Figure 4A-B, treatment with either DHA or DPA did not affect the number of Tfn-Alexa568 puncta after internalization for 5 min (105 % and 106 % of control for DHA and DPA, respectively, p ˃ 0.05). Taken together, these results support that the specific effect of both PUFAs on ligand-induced D2R internalization is not due to an alteration of CME.

**Figure 4:**
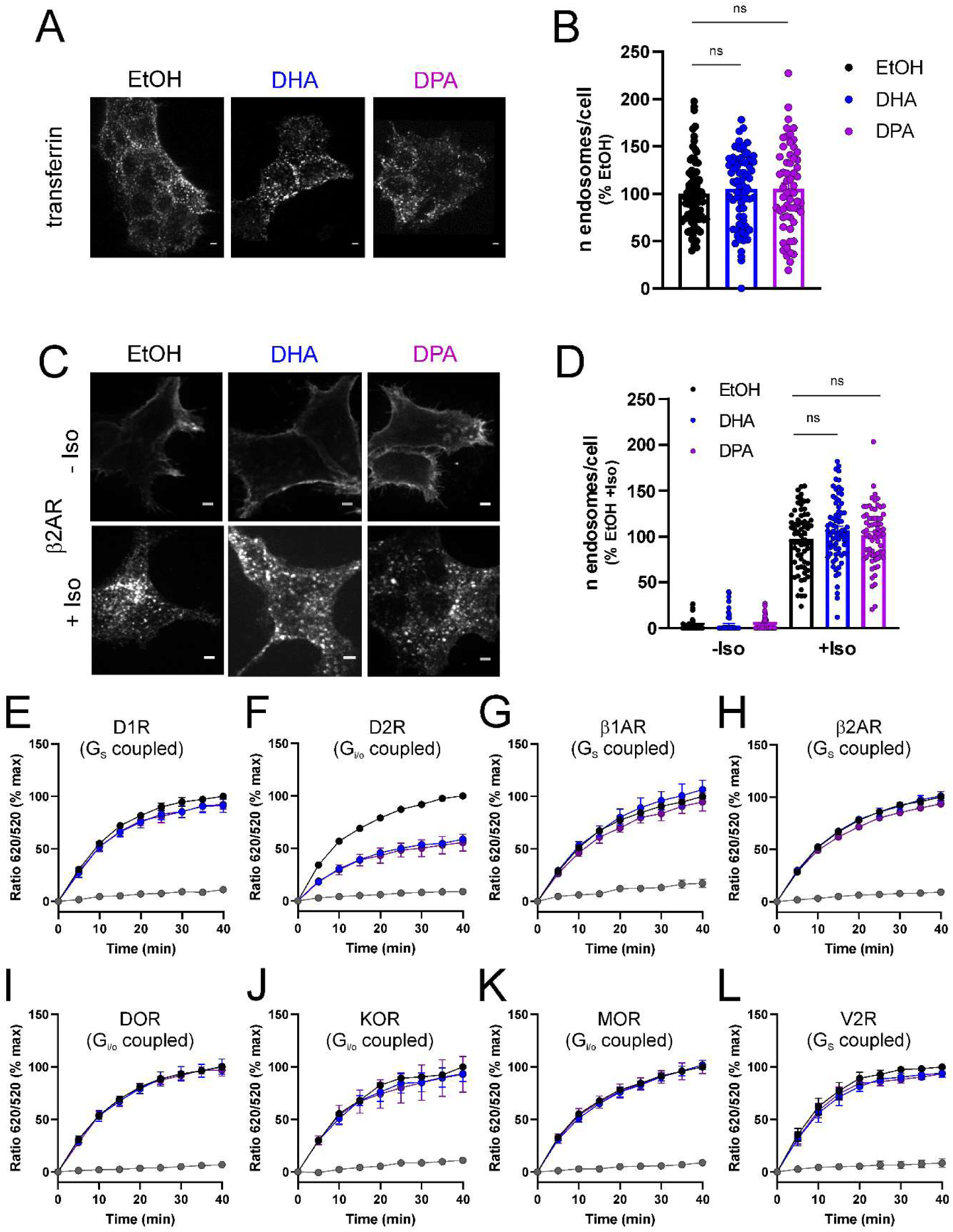
PUFAs do not alter internalization of transferrin or 7 class A GPCRs other than D2R. **A:** Representative confocal microscopy images of control (EtOH) or PUFA-enriched (DHA or DPA) HEK-293 cells incubated with fluorescently labeled transferrin for 5 min at 37 °C. Scale bar 5 μm. **B:** Quantification of the number of transferrin-labelled endosomes relative to control (EtOH); 64-74 cells per condition from 3 independent experiments. One-way ANOVA with Dunnett’s multiple comparisons test: ns *p* ≥ 0.05. **C:** Representative confocal microscopy images of control (EtOH) or PUFA-enriched (DHA or DPA) HEK-293 cells expressing FLAG-β2AR, GRK2 and βarr2, before or after stimulation with 10 µM of isoproterenol (Iso) for 30 min. Scale bar 5 μm. **D:** Quantification of the number of β2AR puncta (endosomes) generated before and after Iso. 64-74 cells per condition collected across 3 independent experiments. One-way ANOVA with Dunnett’s multiple comparisons test: ns *p* ≥ 0.05. **E-L:** DERET assays for internalization of 8 class A GPCRs stimulated (black circles) or not (gray symbols) with selective agonists: SKF 81297 (1 µM) for SNAP-D1R (E); QPL (10 µM) for SNAP-D2R (F); Isoproterenol (10 µM) for SNAP-β1AR (G) and SNAP-β2AR (H); DPDPE (1 µM) for SNAP-DOR (I); U69593 (1 µM) for SNAP-KOR (J); DAMGO (1 µM) for SNAP-MOR (K); AVP (1 µM) for SNAP-V2R (L). Blue and magenta circles represent cells pretreated with DHA or DPA, respectively. The fluorescence ratio 620/520 is normalized to 100 for cells treated with vehicle (EtOH, black circles) at 40 min. Circles and error bars represent mean ± SEM for three independent experiments.

To examine whether the effect of DHA or DPA treatment was specific for the D2R or rather a more general mechanism for class A GPCRs, we examined the internalization of β2 adrenergic receptor (β2AR) following agonist stimulation ^38,39^. As shown in Figure 4C, the FLAG-tagged β2AR was mostly localized at the cell surface in the absence of agonist and internalized after addition of the synthetic agonist isoproterenol (10 µM). In striking difference with the results obtained with the D2R, the number of puncta was similar in control and PUFA-enriched cells (107 % and 101 % of control for DHA and DPA, respectively, p ˃ 0.05) (Figure 4D).

We used again the DERET assay to test the effect of DHA or DPA treatment on various class A receptors: the δ, µ and κ opioid receptors (coupled to G_i/o_ proteins, like D2R) and D1R, β1-and β2ARs as well as vasopressin receptor 2 (V2R) (coupled to G_s_ proteins) (Figure 4E-L). Only D2R showed sensitivity to PUFA treatments (Figure 4E). Altogether, these results suggest that PUFA enrichment specifically affects D2R internalization.

### Alterations of membrane PUFAs in cortical neurons affect D2R endocytosis

Dopamine receptors are highly expressed in neurons. Therefore, we assessed the effect of manipulating PUFA levels on D2R endocytosis in rat primary cortical neurons. We measured with lipidomic analysis the FA levels in primary cultures incubated at 14 DIV for 24 h with 30 µM DHA or DPA. These treatments specifically increased the levels of DHA (from 3.55 ± 0.31 % to 6.13 ± 0.36 %) and DPA (from 0.72 ± 0.03 % to 5.76 ± 0.24 %) upon respective cell incubations (Figure 5A and 5B, respectively). Nevertheless, the total PUFA level is increased only for treatment with DPA (20.53 ± 0.86 % in control, 20.01 ± 1.15 % in DHA treated, 24.36 ± 0.70 % in DPA treated cells, p = 0.0296) (Figure 5C). It should be noted that these values are higher than the ones in HEK-293 cells in all conditions (Figure 1D). We also noted that the proportion of arachidonic acid (20:4 n-6), the most abundant PUFA in brain cells, decreases in DHA treated cells, albeit not significantly (Figure 5D, p = 0.0659), which could reflect a compensation mechanism regulating PUFA levels in neurons. We measured the endocytosis of FLAG-D2R expressed in neurons together with GFP. Incubation of neurons with QPL (10 µM) for 30 min induces a robust endocytosis of D2R. In neurons treated with DHA or DPA, D2R endocytosis is reduced (Figure 5E,F). This result is consistent with the effect of PUFA treatment in HEK-293 cells.

**Figure 5:**
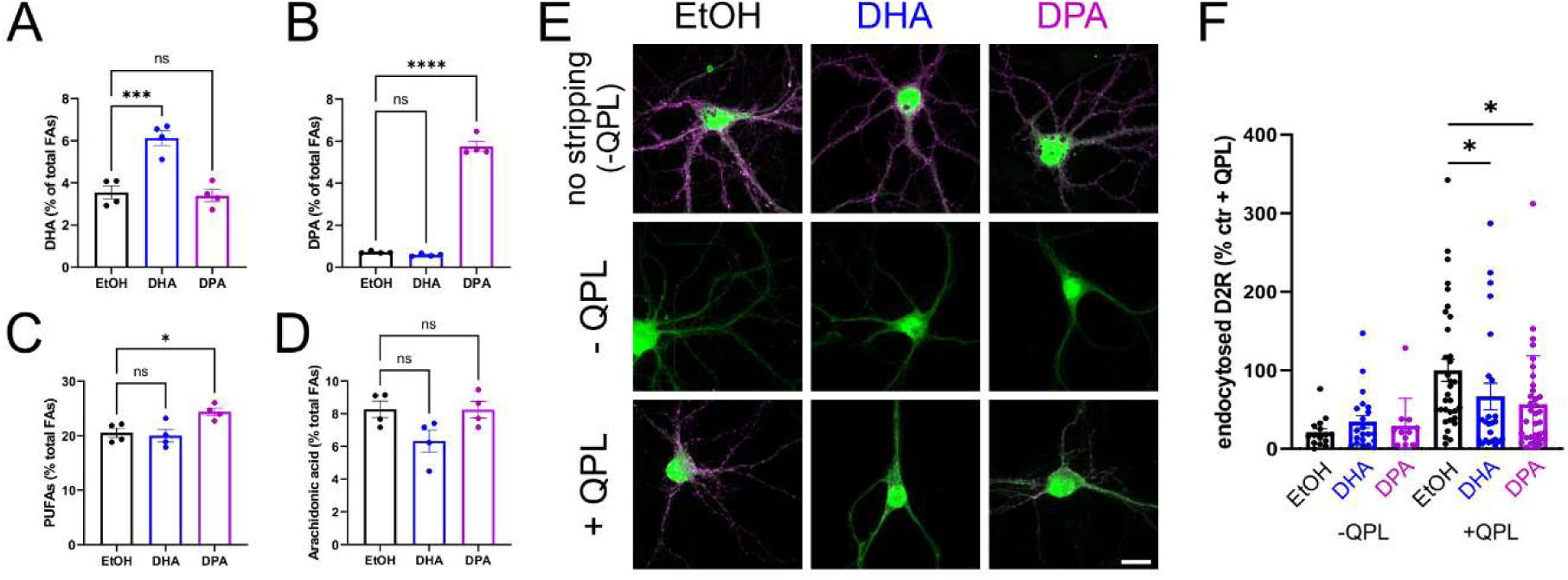
Effect of enrichment in DHA and DPA on D2R endocytosis in neurons in culture. **A-D**, Cellular amount of DHA (A), DPA (B), total PUFAs (C) and arachidonic acid (D) in cortical neurons maintained in culture for 14 days and treated with carrier (0.4 % EtOH, black), DHA (blue) or DPA (magenta) at 30 µM for 24 h. One-way ANOVA followed by Dunnett’s multiple comparison test *** p < 0.001 **** p < 0.0001 ns p > 0.05. **E**, Representative confocal microscopy images of control (EtOH) or PUFA-enriched (DHA or DPA) cortical neurons expressing FLAG-D2R (magenta) and GFP (green), before or after stimulation with either 10 µM QPL for 30 min, and stripped or not. Scale bar: 10 μm. **F**, Quantification of the internalized FLAG-D2R before and after QPL. 15-35 cells per condition collected across 3 independent experiments. One-way ANOVA with Dunnett’s multiple comparisons test: * p < 0.05.

### Membrane PUFAs do not alter D2R clustering but reduce the recruitment of β-arrestin2 and D2R endocytosis

To get insight on the cellular mechanisms underlying blunted D2R ligand-induced internalization under membrane PUFA enrichment, we analyzed the steps of receptor endocytosis following agonist binding: D2R clustering at endocytic pits, recruitment of β-arrestin2 and formation of endocytic vesicles (Figure S3) ^40,41^. Upon agonist stimulation, β-arrestin2 (βarr2) is recruited from the cytoplasm to activated D2R ^31,42^. Once recruited, βarr2 and class A GPCRs cluster at clathrin coated pits with varying degrees ^38,43^. Therefore, we investigated D2R clustering at clathrin-coated pits (CCPs) and β-arr2 recruitment. We imaged living cells expressing D2R tagged with superecliptic pHluorin (SEP), a GFP mutant which is fluorescent at pH 7.4 (extracellular pH) but not at pH 5.5 (endosomal pH), GRK2 and βarr2-mCherry. TIRF microscopy specifically focusses on events happening at the plasma membrane or in its close proximity (∼100 nm). As shown in Figure 6A and Movies S1-3, SEP-D2R signal in control and PUFA-enriched cells is initially homogenous on the plasma membrane, and then clusters following QPL stimulation. We quantitatively assessed this accumulation by measuring the fluorescence intensity at SEP-D2R clusters. Fluorescence quickly increases immediately after agonist application to reach a peak within ∼100 s and slowly decreases thereafter (Figure 6B). The peak of SEP-D2R fluorescence is not significantly different in PUFA-enriched cells compared to control (Figure 6C). We also show in Figure S3B-D that SEP-D2R clusters form at CCPs, labelled with CLC-mCherry, in the same way for all three conditions. βarr2-mCherry is also recruited to SEP-D2R clusters following agonist application (Figure 6D,F). However, treatment with DHA or DPA significantly decreases peak βarr2-mCherry fluorescence (Figure 6F-G). This suggests a blunted βarr2 recruitment at D2R clusters. We obtained similar results with cells treated with dopamine (Figure S4). However, reduced βarr2 recruitment seems at odds with a previous report indicating that cellular enrichment with DHA increases the interaction between D2R and βarr2 after application of QPL, while enrichment with DPA does not affect it ^25^. However, this effect was observed in whole cells after incubation with QPL for 90 minutes, much longer than our total observation time of 10 minutes. To address this apparent discrepancy, we measured with confocal microscopy the recruitment of βarr2-mCherry to intracellular FLAG-D2R clusters after incubation with QPL for 10 and 90 minutes (Figure S5). QPL incubation elicits the formation of intracellular FLAG-D2R clusters at both time points (Figure S5A, B). Consistent with our results showing a reduced number of D2R endosomes at 30 minutes of incubation with QPL (Figure 3), the number of D2R clusters is significantly smaller in cells enriched in DHA or DPA at both 10 and 90 minutes of QPL stimulation (Figure S5B). Nonetheless, while the fluorescence of βarr2-mCherry at FLAG-D2R clusters is similar in all three conditions at 10 minutes incubation, it is significantly higher in cells enriched in DHA at 90 minutes incubation (Figure S5C). Therefore, the recruitment of βarr2-mCherry is differently modulated by the level of DHA and DPA depending on the location (plasma membrane *vs* endosome) and the duration of QPL application. The protein GIPC has also been shown to be involved in D2R endocytosis ^44^. We thus assessed with TIRF microscopy the recruitment of GIPC-mCherry in cells transfected with SEP-D2R, GRK2 and βarr2 following QPL application. We observed only a very weak recruitment of GIPC-mCherry to clusters of SEP-D2R, which was not affected by incubation with PUFAs (Figure S6).

**Figure 6:**
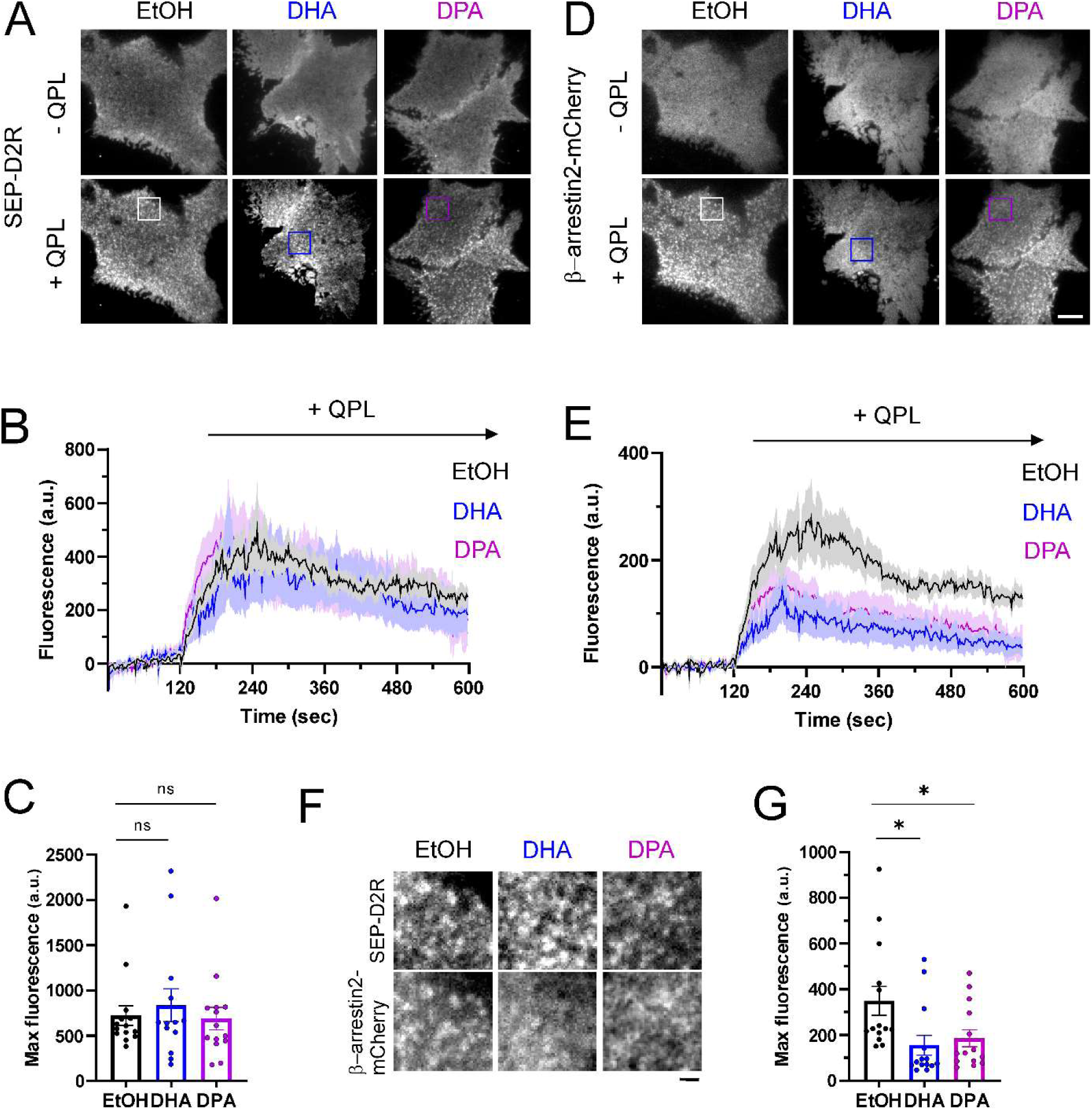
PUFA enrichment does not affect D2R clustering but reduces the recruitment of βarr2 after quinpirole addition. **A, D:** Representative TIRF microscopy images of HEK-293 cells co-expressing SEP-D2R, GRK2 and βarr2-mCherry in control (EtOH) or PUFA-enriched (DHA or DPA) cells, showing the pattern of expression at the plasma membrane for SEP-D2R (A) and βarr2-mCherry (D) before and after stimulation with 10 µM of quinpirole (QPL). Scale bar= 5 µm. See also Movies EV1-3. **B, E:** Mean Fluorescence intensity profiles of SEP-D2R (B) and βarr2-mCherry (E) obtained before and after quinpirole (QPL) addition (120 s after the start of the recording). Data represent average fluorescence values within each cluster minus average fluorescence before agonist addition. **C, G:** Maximum fluorescence intensities of SEP-D2R (C) and βarr2-mCherry (F) measured after QPL addition. Values represent maximum intensity fluorescence minus background fluorescence for 14 cells per condition collected across 3 independent experiments. One-way ANOVA test with Dunnett’s multiple comparisons test; * p < 0.05, ns p ≥ 0.05. **F:** Zoom-in images taken from panels in A and D, as shown by ROIs, depicting SEP-D2R and β-arrestin2-mCherry clusters after quinpirole addition. Scale bar= 1 µm.

To address the direct effect of blunted βarr2 recruitment on endocytic vesicle formation, we tested SEP-D2R endocytosis directly with the pulsed pH (ppH) assay ^45^. In the ppH assay, cells are perfused alternatively every 2 s with buffers at pH 7.4 and 5.5. At pH 5.5, SEP-D2Rs on the plasma membrane are not fluorescent, revealing forming endocytic vesicles (Figure 7A). Endocytic vesicles containing SEP-D2R were thus detected by the ppH assay in images at pH 5.5 at preexisting SEP-D2R clusters, visible at pH 7.4, as shown in an example event (Figure 7B). Following QPL application, SEP-D2R and βarr2-mCherry cluster at the same locations (Figure 7C), like in cells without pH changes (see Figure 6A,E). Moreover, the number of endocytic events detected increases during QPL application and decreases after QPL removal (Figure 7C middle, Figure 7D and Movie S4). While PUFA treatments do not change the number of SEP-D2R clusters detected after QPL application (Figure 7E), it significantly decreases the frequency of endocytic events detected by the ppH protocol (Figure 7F). We then determined the fluorescence of SEP-D2R at the time of vesicle detection. It is affected neither at pH 7.4, reflecting the number of receptors in clusters (Figure 7G), nor at pH 5.5, reflecting the number of receptors in endocytic vesicles (Figure 7H). Moreover, averages of fluorescent traces aligned on vesicle detection (time 0) reveal that the kinetics of endocytic events are not affected by PUFA treatments (Figure 7J, K). Finally, the fluorescence of βarr2-mCherry at SEP-D2R clusters is significantly decreased in both DHA and DPA treated cells, indicating a decreased number of βarr2 molecules recruited (Figure 7I). Moreover, βarr2-mCherry fluorescence was maximal at the time of vesicle detection in all three conditions (Figure 7L).

**Figure 7:**
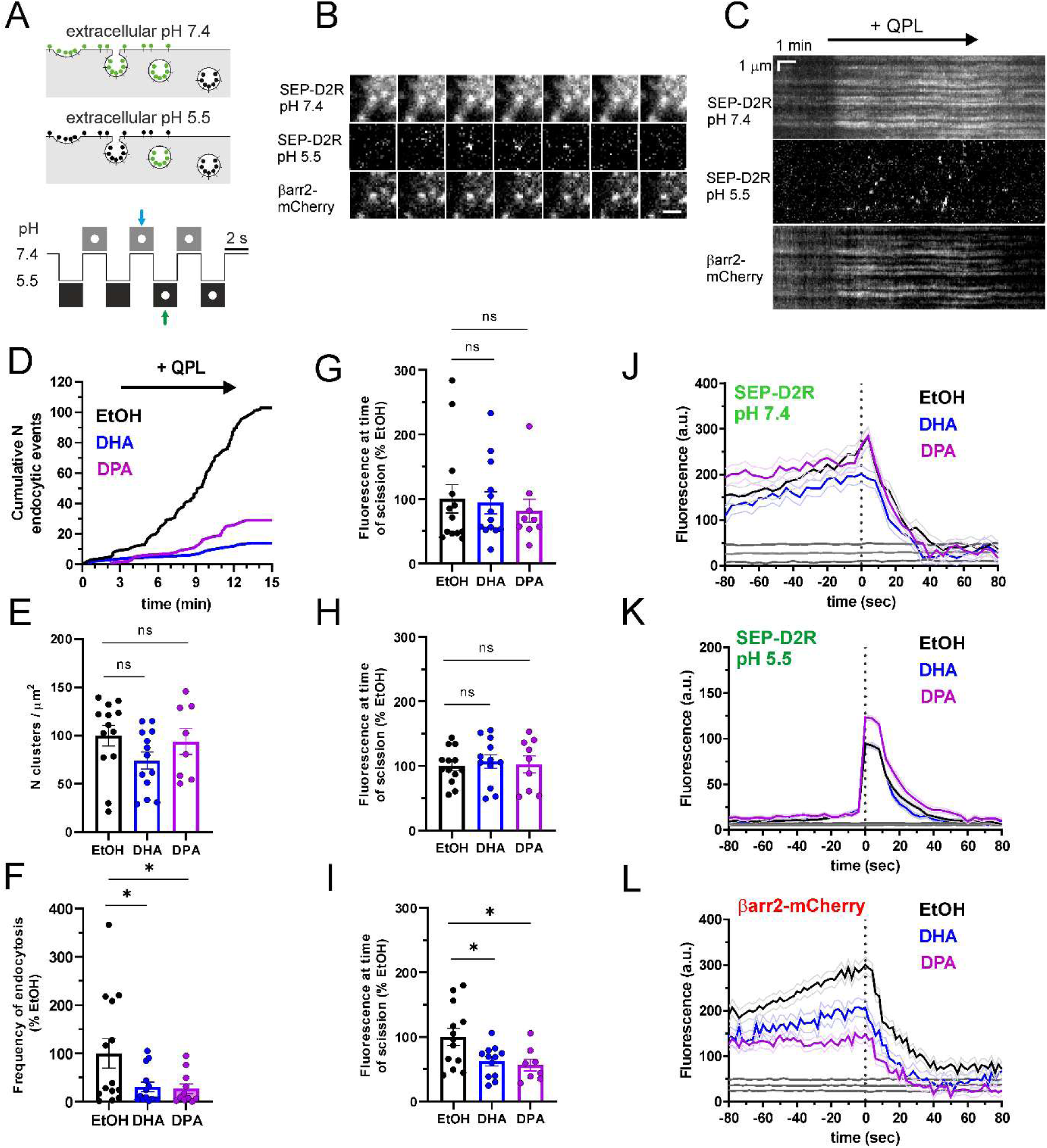
Reduction of endocytic vesicle formation and recruitment of βarr2 at nascent D2R endocytic vesicles in PUFA enriched cells. **A:** Top, experimental procedure: a cell expressing SEP-D2R is bathed in solutions alternating between pH 7.4 and 5.5. At pH 7.4, SEP-D2R on the plasma membrane and in non-acidic vesicles is visible (green lollipops). At pH 5.5, the surface receptors are not fluorescent (black lollipops), and the fluorescence from non-acidic CCVs is isolated. Bottom, principle of vesicle detection: a CCV formed at pH 7.4 (blue arrow) will be visible on the next image at pH 5.5 (green arrow). **B:** Example of a D2R cluster (top row), the corresponding endocytic event (middle row) and the associated β-arrestin2 cluster (bottom row) detected with the ppH protocol. Scale bar: 1 µm. **C:** Kymographs from a representative movie (Movie EV4) of a HEK-293 cells transfected with SEP-D2R, βarr2-mCherry and GRK2 and imaged live with the ppH protocol before (0-120 s), during (121-720 s) and after (701-900 s) application of 10 μM Quinpirole (QPL). **D:** Cumulative number of endocytic events detected with the ppH protocol in control (EtOH) or PUFA-enriched (DHA or DPA) representative HEK-293 cell transfected and imaged as in C. **E:** Number of D2R clusters/cell detected during Quinpirole application in cells transfected as in C, treated with either EtOH, DHA or DPA and imaged live with the ppH protocol. One-way ANOVA with Dunnett’s multiple comparisons test of 13, 13 and 8 cells treated with EtOH, DHA or DPA, respectively; ns p ≥ 0.05 **F:** Number of endocytic events/min/µm^2^ detected during Quinpirole application in cells transfected as in C, treated with either EtOH, DHA or DPA and imaged live with the ppH protocol. One-way ANOVA with Dunnett’s multiple comparisons test of 14, 14 and 11 cells treated with EtOH, DHA or DPA, respectively; ** p < 0.01, * p < 0.05. **G-I:** Average fluorescence intensity of all events in a given cell, at the time of endocytic event detection, for SEP-D2R at pH7.4 (G), SEP-D2R at pH 5.5 (H) and β-arrestin2-mCherry at pH 5.5 (I) obtained from cells transfected and imaged as in C. One-way ANOVA test with Dunnett’s multiple comparisons test for n= 13, 12 and 9 cells treated with EtOH, DHA and DPA, respectively; ** p < 0.01, * p < 0.05, ns p ≥ 0.05. **J-L:** Average fluorescence over time of SEP-D2R at pH 7.4 (J), SEP-D2R at pH 5.5 (K) and β-arrestin2-mCherry at pH 5.5 (L), aligned to the time of vesicle scission (t = 0 s) obtained from terminal events (events for which the SEP-D2R cluster at pH 7.4 disappears within 80 s, defined as in ^45,82^ for the same cells used in G-I. Black lines represent 95% confidence intervals for randomized events to determine the level of enrichment at sites of scission. 424 terminal events from 13 cells (EtOH), 191 terminal events from 12 cells (DHA) and 254 terminal events from 9 cells (DPA).

Finally, we tested again with the ppH assay the specificity of PUFA enrichment on the endocytosis of the constitutive cargo for clathrin-mediated endocytosis, the transferrin receptor tagged with Lime (TfR-Lime) ^46,47^. Consistent with our previous results on Tfn-A568 uptake (Figure 4A, B), the frequency of endocytic events detected with TfR-Lime or their amplitude in cells incubated with DHA or DPA is not significantly different from control cells (Figure S7). Similarly, we tested with the ppH assay the internalization of SEP-β2AR ^48^ and the recruitment of βarr2-mCherry after application of isoproterenol (100 nM) (Movie S5). Consistent with the results on FLAG-β2AR internalization (Figure 4C, D), treatment with DHA or DPA do not affect the frequency nor the amplitude of endocytic events (Figure S8A-D), or the degree of βarr2-mCherry recruitment to endocytic clusters of SEP-B2AR (Figure S8E, F).

### Mutation of Serines 147 and 148 abolishes the PUFA sensitivity of D2R internalization

What could mediate the sensitivity of D2R to PUFA levels? Molecular dynamics simulation of D2R in membranes enriched or not with DHA showed that the conformation of the ICL2 region changes in these two conditions ^25^. In membranes without DHA, ICL2 sticks out of the membrane into the cytoplasm, making serines 147 and 148 accessible to cytosolic proteins. On the other hand, in membranes enriched with DHA, ICL2 is buried into the plasma membrane, with these two serines interacting mostly with lipid phosphatidyl groups. We thus mutated these two serines into alanines and tested its effect on D2R PUFA sensitivity. Importantly, D2R(S147,148) retains the same affinity for QPL, and the increase in QPL affinity induced by enrichment with DHA, but not DPA, observed with WT D2R ^25^ is not affected in mutant receptors (Figure S9A). Moreover, the changes in conformation induced by QPL and the modulation by DHA or DPA ^25^ are not different between WT and mutant D2R (Figure S9B).

We next measured and tested the internalization of the mutated receptor FLAG-D2R(S147,148A) with confocal imaging of receptors labelled with anti-FLAG antibody for 30 min with or without QPL. In control cells (treated with vehicle), D2R(S147,148A) is exported to the plasma membrane and internalized after quinpirole application as much as WT D2R (Figure 8A, B). However, unlike WT D2R, treatment with DHA or DPA does not affect the internalization of D2R(S147,148A) (Figure 8C). This lack of effect was confirmed with the DERET assay (Figure 8D). Moreover, live cell imaging of cells co-transfected with SEP-D2R(S147,148A) and βarr2-mCherry with the ppH assay showed that, during QPL application, treatments with DHA or DPA do not affect the frequency of endocytic events (Figure 8E), the fluorescence of SEP- D2R(S147,148A) clusters pH 7.4 (Figure 8F) and the recruitment of βarr2-mCherry at receptor clusters (Figure 8G). We conclude that mutations of serines 147 and 148 into alanines selectively abolish the sensitivity of D2R endocytosis to PUFA levels in membranes.

**Figure 8:**
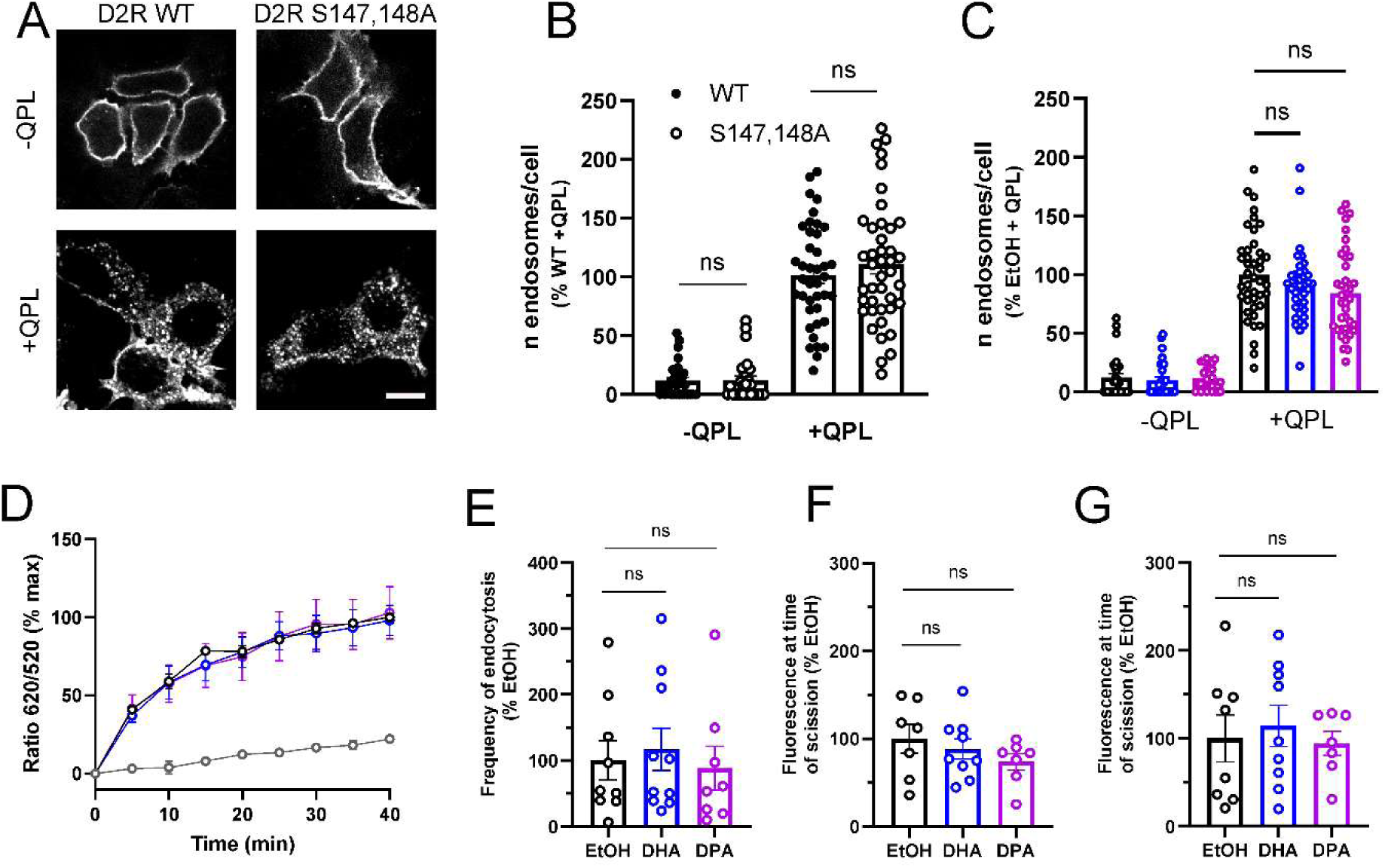
The internalization of the mutant D2R(S147,148A) is not modulated by DHA or DPA. **A**: Representative confocal microscopy images of control (EtOH) HEK-293 cells expressing FLAG tagged D2R or D2R(S147,148A), GRK2 and βarr2, before or after stimulation with 10 µM QPL for 30 min. Scale bar 5 μm. **B**: Quantification of the number of D2R puncta (endosomes) generated before and after QPL stimulation in untreated cells expressing FLAG tagged D2R WT or D2R(S147,148A). 40-42 cells per condition collected across 3 independent experiments. **C**: Quantification of the number of D2R(S147,148A) puncta (endosomes) generated before and after QPL stimulation in control (EtOH) and PUFA enriched cells. 25-42 cells per condition collected across 3 independent experiments; ns *p* ≥ 0.05. **D**: DERET assay for internalization of SNAP- D2R(S147,148A) stimulated (black circles) or not (gray symbols) with QPL. **E:** Frequency of endocytic events detected during QPL application in cells transfected with SEP-D2R(S147,148A), βarr2-mCherry and GRK2, treated with either EtOH, DHA or DPA and imaged live with the ppH protocol (n = 9, 10 and 8, respectively, collected across 3 independent experiments; ns *p* ≥ 0.05). **F, G:** Average fluorescence intensity of all endocytic events in a given cell, at the time of endocytic event detection, for SEP-D2R(S147,148A) at pH 7.4 (F), and β-arrestin2-mCherry (G) obtained from cells transfected and imaged as in E.

## Discussion

Cell surface internalization of GPCRs is a highly regulated and multi-step process in which key protein-protein interactions have been identified ^49,50^. However, little information is available regarding the impact of membrane lipid composition on this mechanism. In the present study, we show that membrane enrichment with either the ω3-PUFA DHA or the ω6-PUFA DPA significantly decreases D2R internalization induced by two full agonists (dopamine and QPL) with three different techniques: DERET, confocal microscopy and TIRF microscopy together with the ppH assay.

One of the most striking results of our study is that altering PUFA levels in HEK-293 cells specifically affects the endocytosis of D2R without altering clathrin mediated endocytosis or the internalization of 7 other class A GPCRs, including G_i/o_ coupled (δ, µ and κ opioid receptors, like D2R) and G_s_ coupled (D1R, β1- and β2AR, V2R) receptors. Membrane enrichment with either DHA or DPA reduces ligand-induced D2R internalization by about half at all concentrations of agonists tested, without affecting its overall internalization kinetics. Specifically, the EC_50_ of QPL-induced internalization is increased, while the EC_50_ of dopamine-induced internalization remains unchanged. On the other hand, the affinities (K_D_) of dopamine or QPL for D2R are increased in membranes enriched with DHA, but not DPA ^25^, suggesting two distinct sites of regulation by PUFAs: one regulating ligand association which is more sensitive to DHA than DPA enrichment, and the other regulating D2R endocytosis, sensitive to both DHA and DPA.

This view is corroborated by molecular dynamics simulations on the D2R which have demonstrated that PUFA-containing phospholipids establish about three times more contacts with the D2R than phospholipids with monounsaturated chains, leading to the formation of a corona around the receptor ^24,25^, as shown also for other class A GPCRs, such as rhodopsin ^51^ and the adenosine A_2A_ receptor ^52^. This preferential interaction could alter the conformation of the D2R in the plasma membrane. Indeed, these simulations have shown that the presence of PUFAs in membrane phospholipids affects mainly two sites on D2R, on one hand the extracellular loops 1 and 2, which contribute to ligand binding ^53,54^, and on the other hand ICL2 on the cytoplasmic side of D2R. The ICL2 of D2R is partially buried in the membrane, especially serines 147 and 148, while it remains fully water accessible in membranes without DHA. The ICL2 of the D2R is rich in polar and charged residues (Figure S10) that would remain exposed to solvent. The PUFA-enriched membrane being more fluid and flexible than the non-enriched one results in a more favourable environment for ICL2 membrane insertion, rendering the process energetically advantageous. ICL2 has been shown to participate in D2R internalization, together with ICL3 ^55^. It is therefore likely that the burying of ICL2 in the plasma membrane under PUFA enrichment could decrease its interaction with endocytic machinery such as β-arrestin. Consistent with this model, we show that mutation of the two consecutive serine residues (Ser 147 and 148), which are located at the ICL2-TM4 junction, does not affect D2R internalization in control, non PUFA-enriched cells, but preserves D2R internalization as well as β-arrestin recruitment in high PUFA conditions. Polar serine residues can establish hydrogen bonds with the phospholipid head groups facilitating its proximity with the membrane. Their replacement by alanines makes the ICL2 significantly more hydrophobic which could change its conformation and membrane embedding properties rendering the double mutant insensitive to PUFAs. Consistent with our two-site model, the S147,148A mutation does not affect the modulation of QPL affinity by DHA.

Interestingly, the two consecutive serines in D2R ICL2 are very rare among class A GPCRs, as shown by sequence alignment (Figure S10 and Supplementary Table 1). None of the 7 tested receptors has it (D1R has two serines located at the junction TM3-ICL2). Among the 200 class A non-orphan/non-opsin GPCRs, only few have consecutive serines in this region and with very different flanking sequences: D3R, nociceptin receptor (NOP), lysophospholipid receptor S1PR5 and melatonin receptor MTR1a. Therefore, very few other class A GPCRs would potentially be sensitive to PUFAs for their internalisation, which could be provided by this double serine signature.

The level of PUFAs in membrane phospholipids could also affect membrane properties directly. For example, PUFA levels affect membrane compartmentalisation in lipid nanodomains, notably the integrity and stability of lipid rafts ^56,57^. However, lipids rafts have been proposed to be signalling platforms for many GPCRs, including the D1R ^58^ or the β2AR ^4,59^, that are insensitive to membrane PUFAs in internalization assays. PUFA levels in membrane lipids could also affect the biophysical properties of the membrane. Indeed, it was shown recently that enrichment of HEK-293 cells in DPA, but not DHA, affects both the elasticity and fluidity of the plasma membrane ^27^. Similarly, incorporation of DHA in retinal pigment epithelial-1 (RPE-1) cells affects their membrane bending rigidity ^60^. Molecular dynamics simulations and biochemical measurements suggest that DHA-containing phospholipids adapt their conformation to membrane curvature *via* their higher flexibility and make the plasma membrane more amenable to deformation by decreasing its bending rigidity ^61^. Consequently, Tfn endocytosis was increased in RPE-1 cells ^60^. Nevertheless, despite a similar level of PUFA enrichment in our protocol (this study) ^25,27^ and in ^60^, we found that PUFA enrichment in HEK 293 cells affects neither Tfn and TfR internalization, nor of other class A GPCRs, arguing against a general effect of PUFA enrichment on endocytosis.

PUFA enrichment could alter receptor interaction with endocytic machinery such as β-arrestin, GRK, or other proteins such as GIPC, a necessary step for GPCR trafficking ^44,56^. We show with live cell TIRF microscopy in HEK-293 cells that the clustering of D2R to CCPs following QPL application is not affected by DHA or DPA enrichment, but that recruitment of β-arrestin2 to CCPs is decreased by half. Because many GPCRs cluster together with β-arrestins to endocytic structures, the long-standing view was that stable, stoichiometric coupling of a GPCR with β-arrestin is a necessary step for its endocytosis. However, activation of β-arrestin by GPCRs can occur through a recently described catalytic pathway which does not require the formation of a stable complex ^43,62^. In this context, transient activation of membrane-associated β-arrestin by interaction with GPCRs drives its association with CCPs ^62^. It is thus possible to decouple D2R recruitment from β-arrestin recruitment to CCPs, resulting in DHA or DPA membrane enrichment only affecting D2R endocytosis and not its clustering at CCPs. Other proteins involved in D2R endocytosis, such as GIPC, could also be involved in this regulation ^44^. However, we detected only a slight recruitment of GIPC to D2R clusters at the plasma membrane, which is not regulated by DHA or DPA.

We found that enrichment in DHA or DPA for 24 h alters D2R endocytosis in HEK-293 cells as well as in primary cortical neurons in culture. Lipidomic analyses indicate that these two cell types have very different levels of DHA, DPA and overall PUFAs. Therefore, relative alterations of PUFA content in membranes are responsible for the alterations in D2R endocytosis. Moreover, the percentages of PUFAs in cortical neurons cultured for 15 days (∼20 %), and DHA in particular (∼4 %), are lower than the ones measured in the brain of adult animals (total PUFAs ∼30 %, DHA ∼15%) ^63,64^. Interestingly, PUFA levels, and DHA in particular, are lower in the embryonic brain and drop within days of culture ^65,66^. Addition of DHA (1.5 µM) in the culture medium at all times maintains the membrane level of PUFA in cultured neurons, and it increases neurite growth and synapse formation ^65,67^. Similar results were obtained in neurons differentiated from human induced pluripotent stem cells, in which PUFA levels can reach up to 40 % with added DHA ^68^. In the present study, we restricted our analysis to effects of DHA or DPA for 24 h at 14 DIV, after the establishment of synaptic network, to avoid confounding effects on neuronal function. Nevertheless, it will be important to assess the effect of chronic DHA or DPA treatment on neuronal function and D2R endocytosis.

In the brain, GPCR internalization following agonist application mediates specific signalling and receptor desensitization ^69–71^. Interestingly, in substantia nigra pars compacta dopaminergic neurons, which strongly express D2Rs, internalization of the receptors in brain slices following agonist application (dopamine or QPL) was shown to be very limited, whereas another GPCR, µ-opioid receptor, was shown to undergo substantial internalization following agonist application in the same neurons ^72^. Moreover, the same study showed that D2R ectopically expressed in locus coeruleus neurons strongly internalize upon agonist stimulation ^72^. Although a number of associated signalling and scaffold proteins could regulate D2R trafficking, the present study raises the possibility that such an effect could be in part due to different PUFA composition between neuronal subtypes.

In conclusion, our results point to a new mechanism by which lower PUFA levels described in several psychiatric disorders could directly participate in their aetiology by the specific regulation of D2R. Moreover, because dietary intake can modify brain PUFA levels ^10^, the observation that PUFAs can directly alter the functionality and activity of the D2R opens up promising avenues for enhancing the efficacy of traditional pharmacological treatments such as antipsychotics for which the D2R is a main target.

## Materials and methods

### HEK-293 cell culture and transfection

HEK-293 cells (ECACC, #12022001) were grown in DMEM supplemented with 10% Fetal Calf Serum and 1% penicillin/streptomycin in a 5% CO_2_ enriched atmosphere at 37°C. Cells were passed twice a week and used between passages 8 and 20. HEK-293 cells were transfected with Lipofectamine®2000 for transient expression of tagged GPCR, βarr2 and GRK2, or TfR, with parameters adapted for the experiment (see below). SNAP-tagged human GPCRs were purchased from Revvity. FLAG-β2AR was described previously ^73^. SEP-D2R was obtained based on SEP-LHR (Lutenizing Hormone Receptor) and FLAG-D2R (short isoform) constructs described previously in ^73^ and ^74^, respectively. SEP was subcloned from SEP-LHR using AgeI and ligated into FLAG-D2R, containing an AgeI restriction site in the FLAG sequence created by site-directed mutagenesis (QuikChange Mutagenesis kit, Agilent) using oligonucleotides detailed in Reagents and Tools table. TfR-Lime and βarr2-mCherry were described previously (Shen et al. 2023) and ^75^ respectively. FLAG-, SEP- and SNAP-tagged D2R(S147,148A), were created by site-directed mutagenesis (QuikChange Mutagenesis kit, Agilent) using oligonucleotides detailed in Reagents and Tools table and using Flag-, SEP- or SNAP-D2R WT as template, respectively.

For diffusion-enhanced resonance energy transfer (DERET) assays, cells were transfected as follows. White polyornithine 96-well plates were coated with poly-L-lysine (PLL, 50 μl of 0.1 mg/ml) for 30 min at 37 °C, then washed twice with 100 μl sterile phosphate buffered saline solution (PBS). After the washing step, a transfection mixture (50 µl/well) composed of 0.5 µl of Lipofectamine®2000, 5 ng of each of the following plasmids (N-terminal SNAP tagged GPCR, GRK2 and βarr2, alone or in combination) and 185 to 195 ng of an empty vector (to reach a total amount of 200 ng of plasmid) were prepared in Opti-MEM reduced serum medium. After 20 min incubation at room temperature, the DNA/lipofectamine mixture was added to each well and followed by 100 µl of HEK-293 cells at a density of 250,000 cells/ml. Cells were grown 24 hours at 37 °C before the addition of PUFAs for 24 h before the DERET measures. For confocal and TIRF microscopy experiments, cells were seeded in a 6 well plate in order to reach approximately 70 % confluence on the day of the transfection. Cells were transfected with a mixture (0.5 ml/well) containing 5 µl of Lipofectamine®2000 and 150 ng of each of the following DNA (the N-terminally FLAG-D2R WT, FLAG- D2R(S147,148A) or FLAG-β2AR, the GRK2 and the βarr2) diluted in Opti-MEM medium and added dropwise to each well. For TIRF microscopy, cells were transfected with either 150 ng of SEP-D2R, WT, SEP-D2R(S147,148A) or SEP-β2AR, βarr2-mCherry and GRK2. For experiments with TfR, only TfR-Lime was transfected into the cells. Transfected cells were grown for 24 h at 37°C before incubation with PUFAs for another 24 h.

### Cell fatty acid treatment and analysis

Fatty acids were dissolved in absolute ethanol under nitrogen at 30 mM and stored in aliquots until use. Cells were incubated with the different FAs as in ^25^. Specifically, aliquots of each FA were diluted in complete medium (DMEM for HEK-293 cells, Neurobasal+B27 for neurons) for a final concentration of 30 µM with a final fraction of 0.1% ethanol (EtOH, vehicle for control cells) for 24 h. Following incubation, cells were rinsed with complete medium to remove excess of fatty acids.

In order to ascertain exogenous fatty acid incorporation (DHA, DPA, BA), whole cell fatty acid analysis was performed as in ^25^. Briefly, control and fatty acid treated cells were randomly analyzed according to ^64^. Total lipids were extracted according to the method developed by ^76^ and transmethylated using 7 % boron trifluoride in methanol following ^77^ protocol. Fatty acid methyl esters were extracted and analyzed by gas chromatography-flame ionization detector (GC-FID, Hewlett Packard HP5890, Palo Alto, CA). Fatty acid methyl esters were identified by comparison with commercial and synthetic standards of saturated, monounsaturated and polyunsaturated FAs (see list in source data) and data analysis were processed using the Galaxie software (Varian). The proportion of each fatty acid was expressed as a percentage of total fatty acids to allow the comparison of lipid composition in different cell culture conditions.

### Neuronal cultures and transfections

Sprague-Dawley rats (Janvier Labs) of either sex were euthanized in accordance with the European 2010/63/EU directive, Annex IV Methods of killing animals. All experiments were performed in accordance with the European guidelines for the care and use of laboratory animals, and the guidelines issued by the University of Bordeaux animal experimental committee (CE50; Animal facilities authorizations A3306940, A33063941). Cortical neurons were obtained from E18 rat embryos. For lipidomic analysis, we plated 7.10^6^ dissociated cortical cells on 75 cm^2^ flasks and cultured them in complete neuronal medium, composed of Neurobasal™ Plus medium supplemented with 2 mM L-glutamine and B-27™ Plus, renewed by third twice a week for 14 DIV. At 6 DIV, we added cytosine arabinoside (2 μM) to prevent glial proliferation. For transfection and confocal imaging, we plated dissociated cortical cells at a density of 200,000 cells per 60 mm dish onto poly-L-lysine-coated coverslips. After 2 hours, the coverslips were placed in a ’sandwich configuration’ on dishes containing glial cells. Cultures were maintained at 36.5°C in a 5% CO2 atmosphere in complete neuronal medium. To prevent glial cell proliferation, we added cytosine arabinoside (2 μM) to the media after 3 DIV. The neurons were transfected at 6 or 7 DIV using a calcium phosphate technique. In brief, prior to transfection, the culture medium was removed and replaced with fresh medium. The calcium phosphate precipitate was formed by mixing 4 μg of FLAG-D2R or FLAG-D2R(S147, 148A), 0.5μg GFP, with nuclease-free water (per 4 12mm coverslip), 250 mM CaCl2, and equal volume of 2x BES buffered saline (BBS) (100 μL). The precipitate was incubated for 15 minutes at RT in the dark and added dropwise to the neurons and incubated for 20 minutes at 37°C, during which a fine sandy precipitate covered the cells. The neurons were then washed twice in HBSS and transferred back to their original culture medium. Transfected cells were grown till DIV15-16 at 37°C, with partial refreshment of the culture medium twice per week. Neurons were used for immunocytochemistry and confocal imaging at DIV15-16, with prior incubation of PUFAs for 24 h (see below for detailed protocol of incubation).

### Cell viability assay

Cell viability was measured using Resazurin Fluorimetric Cell Viability Assay. One day after treatment, control (ethanol) and fatty acid enriched cells were washed gently twice with pre-warmed PBS, and then a mixture of resazurin and DMEM in a ratio 1:10 was added into each well. After 3 h incubation at 37 °C, cell viability was monitored by measuring fluorescence with λ_ex_ at 540 nm and λ_em_ at 590 nm in a Tecan infinite M1000 Pro reader. The fluorescent signal generated in each well is directly proportional to the number of living cells in the sample. The percentage of survival of fatty acid treated cells was calculated relative to control (EtOH) treated cells.

### DERET assay and data analysis

Internalization assay was performed in white 96-well culture cell plates, using HEK-293 cells transfected with SNAP-D2R, GRK2, βarr2 and incubated with fatty acids, as described above. Cell culture medium was removed 24 h after PUFA treatment and 100 nM of SNAP-Lumi4-Tb (the donor fluorophore) previously diluted in Tag-lite buffer was added (50 μl/well) and further incubated for 1h at 16 °C to prevent/minimize receptor internalization. Excess SNAP-Lumi4-Tb was removed by washing each well four times with 100 μl of Tag-lite labeling buffer.

Membrane expression levels of the receptor were determined by measuring the fluorescence intensity of D2R-SNAP-Lumi4-Tb at 620 nm. Internalization experiments were performed by incubating cells with Tag-lite labeling buffer, either alone or containing one of the D2R ligands (Dopamine, Quinpirole, Aripiprazole, Haloperidol) in the presence of fluorescein (fluorescence acceptor). Typically, in plates containing SNAP-Lumi4-Tb-labeled cells, 90 μl of 25 μM fluorescein was added followed by the addition of 10 µl of ligand to reach a final concentration in the well ranging from 10^-14^ to 10^-4^ M. Fluorescence measurements were performed in a Tecan Infinite M1000 Pro plate reader, at 37°C right after the treatment described above.

For the experiments assessing D2R internalization kinetics, cells were stimulated with 10 μM of ligand. For dose response experiments, signals emitted by the donor (Lumi4-Tb) at 620 nm and by the acceptor (fluorescein) at 520 nm were collected at the indicated time points using time-resolved settings for the donor (1500 μs delay, 1500 μs reading time) and acceptor (150 μs delay, 400 μs reading time). The fluorescence intensity ratio 620/520 (R) was obtained by dividing the donor fluorescence signal (measured at 620 nm) by the acceptor fluorescence signal (measured at 520 nm) at a chosen time.

Data are expressed as percent of maximal internalization [% of max. internalization (620/520 nm)] using the following equation: [(R_t_ – R_min_)/ (R_max_ – R_min_)] × 100 where R_t_ corresponds to the ratio observed at a chosen time, R_min_ and R_max_ correspond, respectively, to the minimum and the maximum fluorescence ratios. Data expressed in dose-response curves were fitted using non-linear regression dose-response and the following equation: Y=Bottom + X*(Top-Bottom) / (EC_50_ + X), where X and Y represent the ligand concentration (log scale) and the maximum internalization (in percentage) allowing to obtain both the maximal effect (Emax) and half maximal effective concentration (EC_50_) values.

### Confocal microscopy imaging and data analysis

For HEK-293 cells, 24 h before confocal imaging, glass slides were placed in a 12-well plate and coated with PLL (250 μl of 0.1 mg/ml) for 30 min at 37 °C. After coating, HEK-293 cells transiently expressing the FLAG-D2R, GRK2 and βarr2 (or FLAG-β2AR, GRK2 and βarr2) were seeded in complete DMEM containing 30 µM of PUFA or 0.1 % of EtOH. Cells were kept at 37 °C, 5% CO_2_ for 24 hours to allow PUFA incorporation into cells. Cells were rinsed with medium and incubated with M1 anti FLAG antibody (1:500) dissolved in complete medium for 10 min at 37° C and then left untreated or stimulated with ligands (dopamine or QPL for D2R, isoproterenol for β2AR) at 10 µM for 30 min. Cells were then washed with PBS (pH 7.4 containing 0.9 mM of Ca^2+^) and fixed with 4% (v/v) paraformaldehyde diluted in Ca^2+^ enriched PBS for 10 min at room temperature. Fixed cells were permeabilized with 0.2% (v/v) Triton-X for 10 min and incubated with Alexa Fluor-555 anti-mouse antibody (1:1000) for 30 min at room temperature in the dark. Cells were washed 3 times with PBS and the coverslips were mounted on glass slides using Fluoromount-G.

Cortical neurons expressing FLAG-D2R and GFP on individual coverslips were incubated with M1 anti-FLAG monoclonal antibody (1:500) at 37 °C for 10 min. Neurons were then left untreated or stimulated with QPL at 10 µM for 10 min. Cells were then washed with PBS (pH 7.4 containing 0.9 mM of Ca^2+^). To strip anti-FLAG antibody from residual surface receptors (those not internalized by agonist), cells were washed three times in PBS containing 0.04% EDTA and lacking Ca^2+^ (M1 interaction is Ca^2+^ dependent). Cells were fixed with 4% (v/v) paraformaldehyde diluted in Ca^2+^ enriched PBS for 10 min at room temperature. Fixed cells were permeabilized with 0.2% (v/v) Triton X-100 for 5 min and incubated with 10% Bovine Serum Albumin (BSA) for 30 min, followed by incubation with Alexa Fluor-555 anti-mouse antibody (1:1000) for 45 min at 37 °C. Cells were washed 3 times with PBS and the coverslips were mounted on glass slides using Fluoromount-G.

Images were acquired with a spinning-disk confocal microscope (Leica DMI 6000B) controlled with MetaMorph (Molecular Devices), equipped with an EMCCD evolve camera and 488 and 561 nm LASER diodes. Images were analyzed using MetaMorph software and Matlab 2018a (MathWorks). Z-stack of twelve images separated with a fixed step size of 0.5 μm were acquired with a 63×/1.4 NA oil immersion objective. The maximal projections of the z-stacks were used to quantify the internalized receptors. A mask was traced around each cell (cell outline for HEK-293 cells, GFP mask for neurons) as well as a background region using MetaMorph. For HEK-293 cells, the cell mask was eroded of 3 pixels (360 nm) to exclude the plasma membrane region containing receptors on the plasma membrane. Generated clusters of internalized receptors (FLAG-D2R or FLAG-β2AR) were detected by wavelet segmentation using custom written Matlab programs. For each cell, the number of clusters per mask (for HEK-293 cells) or the number * (cluster fluorescence) per mask (for neurons) was computed using a homemade Matlab script. These numbers were normalized to 100 for the control condition (cells grown in absence of PUFAs and incubated with receptor agonists) for each of the 3 independent experiments performed.

### Transferrin uptake assay

HEK-293 cells were seeded on PLL coated glass coverslips and incubated with PUFAs as described above. After 24 h, cells were serum starved in pre-warmed DMEM for 30 min and then incubated for 15 min in cold DMEM containing 10 µg/ml Alexa Fluor 568-conjugated transferrin (Tfn-A568). Cells were incubated 5 min at 37 °C, allowing Tfn-A568 internalization, and washed in cold DMEM then in cold glycine buffer (100 mM NaCl, 50 mM glycine, pH 3) for 2 min to remove Tfn-A568 bound to non-internalized TfR (stripping). Stripped cells were then fixed as explained above and kept at 4 °C until imaging. Fixed cell imaging and cluster quantification were performed in the same manner as described above for the GPCRs.

### TIRF microscopy imaging and data analysis

Cells transiently expressing SEP tagged D2R, GRK2 and βarr2-mCherry were plated on PLL coated glass coverslips, grown in complete DMEM and then enriched with PUFAs as described above. 48 h after transfection and 24 h after PUFA enrichment, live cells were washed 2 times with FluoroBrite™ and imaged for 10 min in the same medium. The indicated D2R agonist was added at time 120 s, for experiments shown in time-courses. Live TIRF imaging was performed at 37°C in a temperature-controlled chamber under constant flow on a motorized Olympus IX83 inverted microscope outfitted with TIRF illuminator (Ilas2, Gataca Systems) with fiber coupled Laser sources (Cobolt Lasers 06-DPL 473 nm, 100 mW and 06-MLD 561 nm, 50 mW). Images were obtained with an Olympus TIRF objective (150X, 1.45 NA, oil immersion). Dual channel live acquisition was performed using 2 EMCCD cameras (QuantEM Model 512SC, Princeton Instruments) controlled by Metamorph 7.10 software (Molecular Devices). The gain of the EMCCD cameras was kept constant at 600, binning at 1 × 1, readout speed 10 MHz. Time lapse image sequences were acquired for up to 20 minutes at 0.5 Hz with an exposure time of 100 ms. Images of fluorescent beads (Tetraspeck microspheres, 0.2 µm) were acquired in every experiment as described previously ^45^. Micro bead images were used to determine the precise parameters required for alignment of the two cameras; these parameters were then applied to all the images acquired on that day.

To quantify SEP-D2R and βarr2-mCherry fluorescence at clusters, the 11 images taken 100-120 s after addition of agonist, which corresponds to full clustering of SEP-D2R (see Figure 5B), were averaged. Clusters were detected by wavelet segmentation using custom written Matlab program. The movies were corrected for XY drift. Fluorescence values measured in the cluster regions were computed. The fluorescence values of the 60 frames before agonist addition were averaged and subtracted. The regions determined for SEP-D2R were transferred to the βarr2-mCherry channel with correction as described in the previous paragraph to compute βarr2-mCherry fluorescence values.

### Pulsed pH (ppH) protocol

Cells transfected (with either SEP-D2R, GRK2 and βarr2-mCherry; SEP-D2R(S147,148A), GRK2 and βarr2-mCherry; SEP-β2AR, GRK2 and βarr2-mCherry; TfR-Lime) and enriched in PUFAs as described in the paragraph above were perfused with HEPES buffered saline solution (HBS) at 37 °C. HBS contained, in mM: 135 NaCl, 5 KCl, 0.4 MgCl_2_, 1.8 CaCl_2_, 1 D-glucose and 20 HEPES, and was adjusted to pH 7.4 and osmolarity of the medium cells were growing in. MES buffered saline solution (MBS) was prepared similarly by replacing HEPES with MES and adjusting the pH to 5.5 using NaOH. All salts were from Sigma Aldrich. HBS and MBS were perfused locally around the recorded cell using a 2-way borosilicate glass pipette as described in ^45^. Movies were acquired at 0.5 Hz for a total of 900 s (GPCR) or 300 s (TfR). For experiments with SEP-D2R or SEP-β2AR, cells were imaged for 120 s, then agonist was added in both HBS and MBS channels at a final concentration of 10 µM (quinpirole) or 100 nM (isoproterenol) for 600 s, then was washed out for the remaining 180 s. For experiments with TfR-Lime cells were imaged for 300 s.

Detection of endocytic events and their analyses were conducted using custom made Matlab scripts previously described ^45,78,79^, apart from kymographs which were obtained using ImageJ. Endocytic events were automatically detected, defined as sudden punctate fluorescence increase appearing in pH 5.5 images, visible for more than 3 frames (i.e. 8 seconds) and appeared at the same location as a pre-existing fluorescence cluster detectable in pH 7.4 images. Event frequency was expressed per cell surface area measured on the cell mask. Fluorescence quantification of events was performed as in ^45^. In short, each value is calculated as the mean intensity in a 2-pixel (206 nm) radius circle centered on the detection to which the local background intensity is subtracted (the local background is taken as the 20th to 80th percentile of fluorescence in an annulus of 5 to 2 pixel outer and inner radii centered on the detection). The frame of maximum intensity is used as the reference (time 0) for aligning and averaging all traces. Before (resp. after) tracking of an object the fluorescence is measured at the location of the first (resp. last) frame with the object tracked. Two color alignment was performed using an image of beads (TetraSpeck microspheres, ThermoFisher) taken before the experiment.

### Plasmon waveguide resonance (PWR) measures

PWR experiments were conducted using a homemade instrument equipped with a He–Ne laser (λ = 632 nm) that produces linearly polarized light at 45°. This configuration enables the simultaneous acquisition of both *p*-polarized (parallel to the incident light) and *s*-polarized (perpendicular to the incident light) signals within a single angular scan. The sensor consists of a right-angle (90°) prism whose hypotenuse surface is coated with a 50 nm silver layer and a 460 nm silica overlayer. The prism is coupled to a Teflon block containing the sample cell. The entire assembly is mounted on a rotating table connected to a motion controller (Newport XPS; ≤ 1 mdeg resolution). Reflected light is measured as a function of the incident light angle by a photodiode (Hamamatsu) ^80^. All experiments were performed at a controlled room temperature of approximately 23 °C.

The protocol for the adhesion of cell membrane fragments onto the silica surface of the sensor has been described previously ^81^. Briefly, the silica surface was rinsed with ethanol, followed by cleaning and activation in a plasma cleaner for 2 min (Diener, Bielefeld, Germany). Subsequently, the surface was incubated with a solution of poly-L-lysine (PLL, 0.1 mg/mL) for 40 min and then thoroughly washed with phosphate-buffered saline (PBS). Cells grown to less than 50% confluence were rinsed with PBS and then with deionized water to induce osmotic swelling. Immediately afterward, the sensor was placed in direct contact with the swollen cells and a slight pressure was applied for approximately 2 min to promote mechanical rupture of the cells and facilitate the transfer of plasma membrane fragments onto the silica surface. The sensor was rinsed with PBS to remove residual cellular debris, mounted with the PWR cell chamber (total volume 250 μL) and filled with buffer. PWR measurements were conducted immediately thereafter.

Ligand-binding experiments were performed by incrementally adding quinpirole aliquots (dissolved in PBS buffer) to the PWR cell chamber (assembled with the PWR sensor with immobilized cell membrane fragments) while continuously monitoring the corresponding PWR spectral shifts for both polarizations. The initial ligand concentration was selected to be approximately one order of magnitude below the reported dissociation constant (KD) for the compound. Following each incremental addition, the system was allowed to reach equilibrium prior to recording the PWR response. KD values were determined by plotting the angular position of the PWR resonance minimum, reflecting ligand-induced receptor conformational changes, as a function of ligand concentration in the cell. The resulting binding isotherms were fitted to a hyperbolic function describing a 1:1 ligand–receptor interaction using GraphPad Prism (GraphPad Software). Data obtained from *p*- and *s*-polarized measurements were analyzed independently, and the mean of the two KD values was reported. The magnitude of the spectral shifts observed at ligand-saturating conditions, corresponding to the maximum binding response (Bmax), was also quantified.

### Statistical analysis

Data are presented as mean ± SEM (standard error of the mean). Statistical significance was determined using GraphPad Prism 8.3.0 (GraphPad, La Jolla, CA, USA). Normally distributed data sets were analyzed using one-way ANOVA or two-way ANOVA followed by Dunnett’s or Sidak’s post-test were used when comparing more than two groups or at least two groups under multiple conditions, respectively. Non-normally distributed data sets were analysed using Kruskal-Wallis test followed by Dunn’s post-test. Differences were considered significant at p < 0.05. See Table 2 for values of all statistical tests performed in this study.

## Acknowledgements

The authors are grateful to Prof Jonathan Javitch for providing the FLAG-D2R construct, Prof Aylin Hanyaloglu for the FLAG-β2AR construct, Dr Philippe Marin for the GRK2 construct, Dr Stefano Marullo for the β-arrestin2-mCherry construct and Dr Daniel Choquet for the GIPC-GFP construct. Imaging experiments involving spinning disk confocal microscopy were carried out at the Bordeaux Imaging Center, a service unit of the CNRS-INSERM and Bordeaux University, member of the national infrastructure France BioImaging supported by the French National Research Agency (ANR-10-INBS-04). We thank Magali Mondin for help with confocal microscopy. We thank MetaToul (Toulouse metabolomics & fluxomics facilities, https://mth-metatoul.com/) which is part of the French National Infrastructure for Metabolomics and Fluxomics MetaboHUB-ANR-11-INBS-0010). Figures S1A and S3A were created with BioRender®. The work was supported by the Agence Nationale de la Recherche (polyFADO ANR-21-CE44-0019 to TD, PT, DP and IA; LocalEndoProbes ANR-14-CE16-0012 and DopamineHub ANR-19-CE16-0003 to DP; FrontoFat ANR-20-CE14-0020 to PT), the Fondation Recherche Medicale team grants to PT and DP, University of Bordeaux’s IdEx “Investments for the future” program/GPR BRAIN_2030 to PT and DP. SS is supported by a Sir Henry Wellcome postdoctoral fellowship (218650/Z/19/Z). RB obtained a PhD fellowship from the French Ministère de l’Enseignement Supérieur et de la Recherche.

## Author contributions

SS, RB, TD, PT, DP, IA designed the study. SS, RB, ML, AAM, VDP, JH, AG, DP, IA performed the experiments and analyses. TD, PT, DP, IA provided funding. SS, RB, PT, DP, IA wrote the initial versions of the manuscript. All authors edited the manuscript.

