## Supplementary Figures S1-10 for "Membrane lipid poly-unsaturation selectively affects dopamine D2 receptor internalization"

### Supplementary Figures and Movies

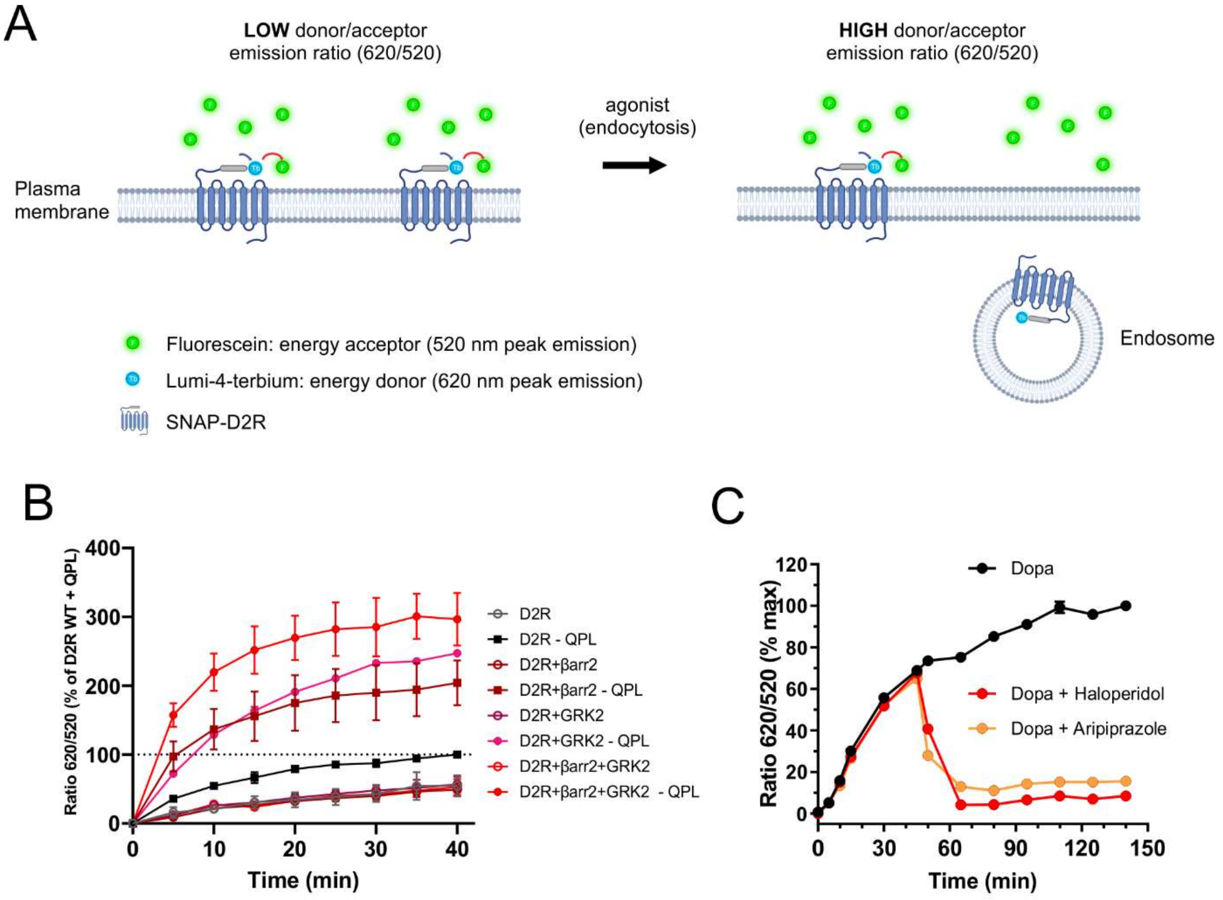


**Figure S1: Validation of the DERET assay to measure D2R internalization after agonist stimulation**. **A**, Principle of DERET assay. In absence of its agonist, the D2R covalently labeled with cell-impermeable SNAP-Lumi4®-Tb (energy donor, blue) is at the cell surface. Addition of excess of fluorescein (energy acceptor, green dots) in the extracellular medium leads to efficient energy transfer (red arrows) resulting in a low DERET ratio (620/520 nm). Following agonist addition (yellow dots), D2R internalizes which causes a significant reduction of energy transfer to the acceptor resulting in higher DERET ratio. **B**, Real-time internalization of SNAP-D2R following application of QPL (10 µM) (filled squares) or not (hollow circles) at time 0 in cells transfected with SNAP-D2R alone (black) or in combination with β-arrestin2 (brown), GRK2 (pink), or both (red). **C**, Real-time internalization of SNAP-D2R following Dopamine (Dopa) addition alone (10 µM, at t0) or upon addition at t = 50 min of the antagonist Haloperidol (100 µM) or the partial agonist Aripiprazole (100 µM). In B and C, Percentages of fluorescence ratio R (620/520 nm) are plotted as a function of time.

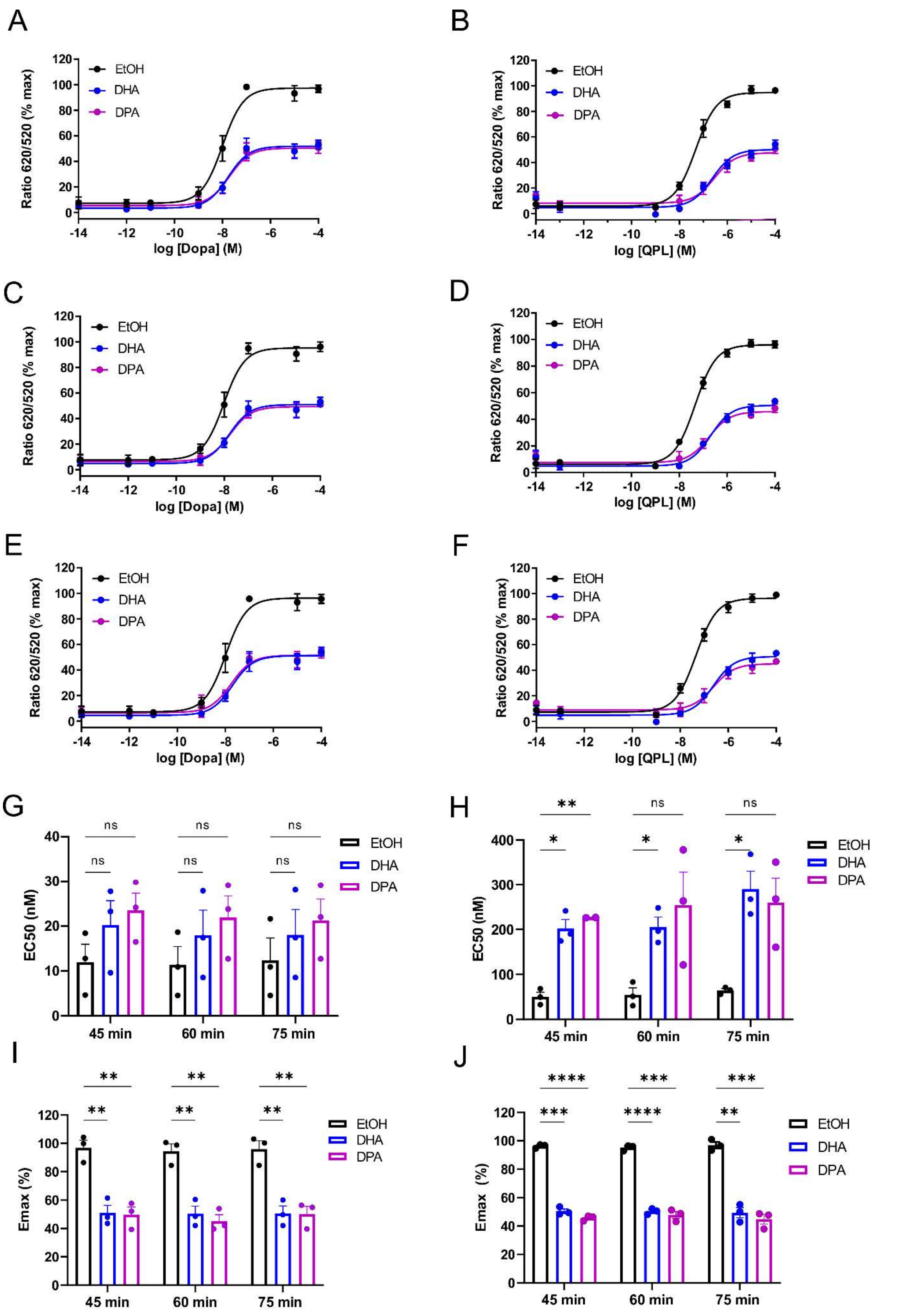


**Figure S2: Impact of PUFAs on D2R induced internalization investigated by DERET assay A-F**: Dose–response curves of D2R induced internalization. Control and PUFA-enriched

(DHA and DPA) HEK cells expressing SNAP-D2R were incubated in the presence of an increased concentration of either Dopamine (Dopa) or Quinpirole (QPL) for 45 min (A, B), 60 min (C, D), 75 min (E, F) at 37°C. Data were fitted using non-linear regression dose–response log [ligand] versus response with three parameters. **G**, **H**: EC50 values calculated from dose-response curves obtained at later time points (45, 60, 75 min) following stimulation with either Dopamine (G) or Quinpirole (H). **I**, **J**: Emax values measured from dose-response curves obtained at later time points (45, 60, 75 min) following stimulation with either Dopamine (I) or Quinpirole (J). Two-way ANOVA with Dunnett's multiple comparisons test of n=3 independent experiments carried out in triplicates; **** $p < 0.0001$, ** $p < 0.01$, * $p < 0.05$, ns $p \geq 0.05$.

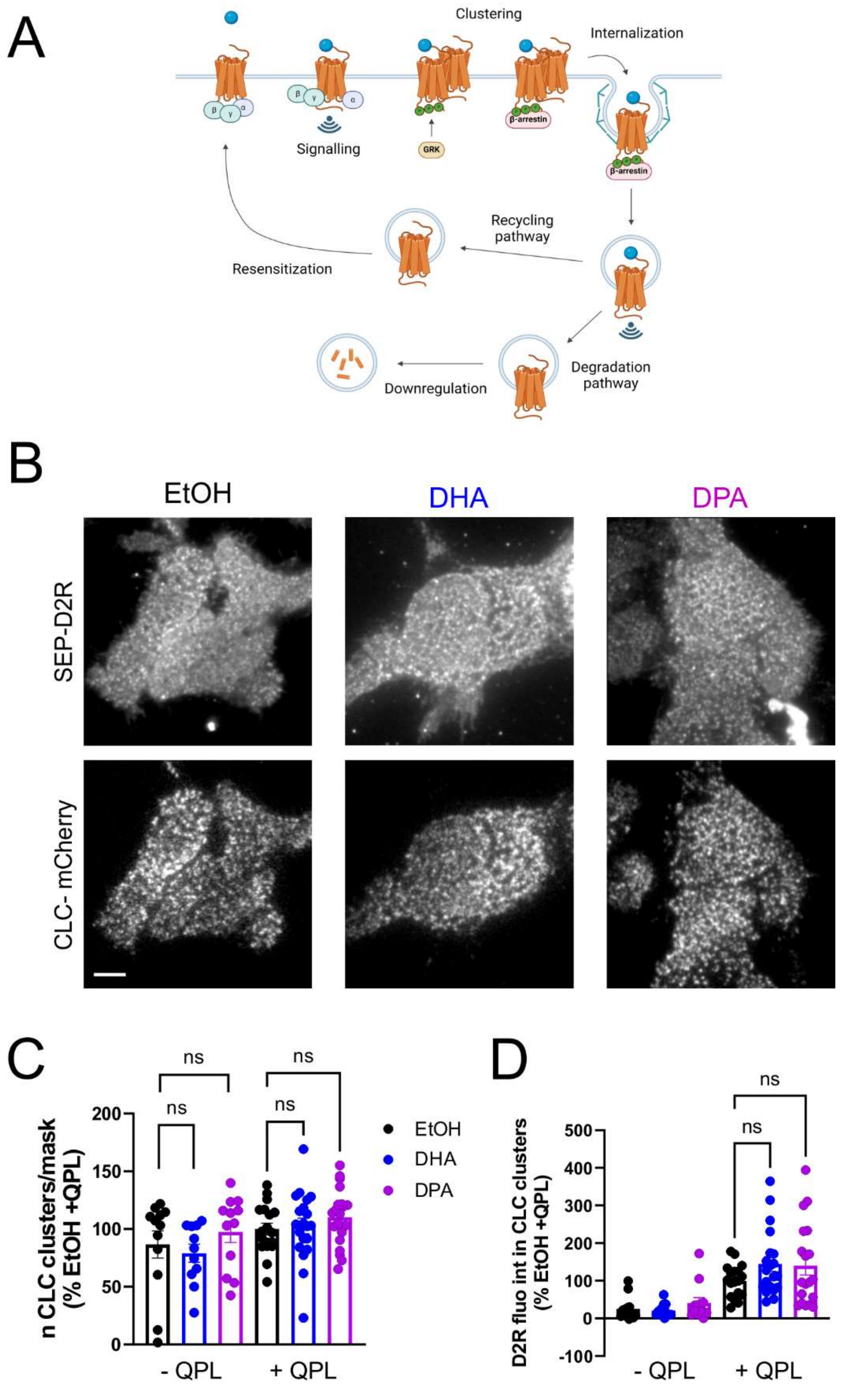


**Figure S3: PUFA treatments do not affect clustering of D2R to CCPs. A**, Scheme of the key steps and proteins involved in D2R internalization, which takes place at CCPs. **B**, Representative images of HEK-293 cells transfected with SEP-D2R, CLC-mCherry,β-arr2 and GRK2, enriched in either ethanol (EtOH), DHA or DPA, treated with QPL (10 µM) for 10 minutes and imaged live by TIRF microscopy. Scale bar = 5 µm. **C**, Quantification of the number of CCPs per cell, from cells as in B n = 12-20 cells/condition collected across 3 independent experiments. One-way ANOVA followed by Dunnett's multiple comparison test. ns $p>0.05$. **D**, Quantification of D2R fluorescence in CCPs relative to D2R fluorescence outside

CCPs, before (-QPL) and after stimulation with QPL (+QPL), from cells as in B n=12-20 cells/condition collected across 3 independent experiments. One-way ANOVA followed by Dunnett's multiple comparison test. ns $p \geq 0.05$.

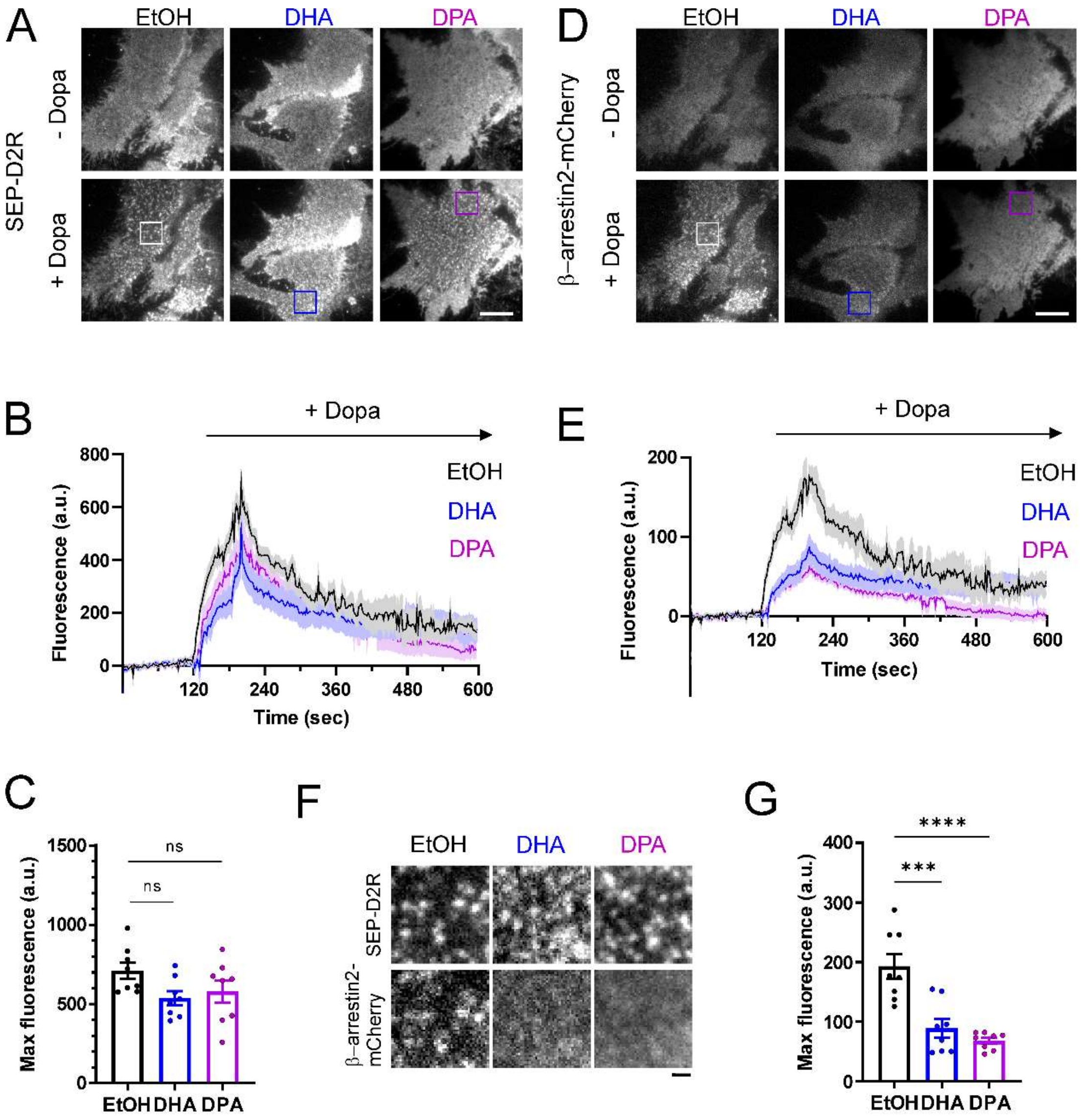


**Figure S4: PUFA enrichment does not affect D2R clustering but impairs the recruitment of β-arr2 upon Dopamine addition. A, D**: Representative TIRF microscopy images of HEK293 cells co-expressing SEP-D2R, GRK2 and β-arrestin2-mCherry enriched in either ethanol (EtOH), DHA or DPA, showing the pattern of expression at the plasma membrane for SEP-D2R (A) and β-arrestin2-mCherry (D) before and after stimulation with 10 µM of Dopamine (Dopa). Scale bar 5 µm. **B, E:** Mean Fluorescence intensity profiles of SEP-D2R (B) and β-arrestin 2-mCherry (E) obtained before and after Dopamine (Dopa) addition at t=120 s. Data represent average fluorescence values within an ROI drawn around each cluster minus background fluorescence (= fluorescence measured in the ROI during the 60 frames before agonist addition was averaged and subtracted from fluorescence values at each frame). **C, G:** Maximum fluorescence intensities of SEP-D2R (C) and β-arrestin2-mCherry (G) measured after Dopamine addition. Values represent maximum intensity fluorescence minus background fluorescence in each ROI for n= 8 cells per condition collected across 2 independent experiments. One-way ANOVA test with Dunnett's multiple comparisons test; *** $p < 0.005$, ns $p \geq 0.05$. **F:** Zoom-in images taken from panels in A and D, as shown by ROIs, depicting SEP-D2R and β-arrestin2-mCherry clusters after Dopamine addition. Scale bar= 1 µm.

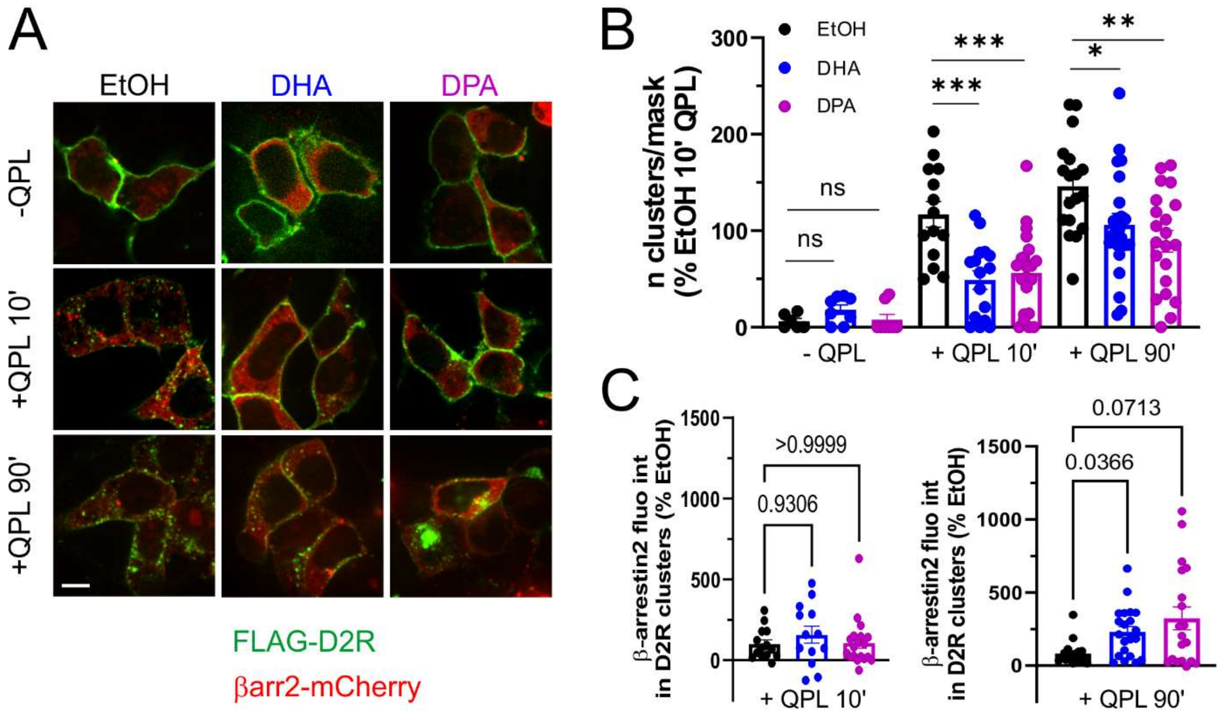


**Figure S5: DHA treatment increases recruitment of βarr2 to D2R intracellular clusters at 90 minutes of QPL stimulation.** **A**, Representative confocal microscopy images of HEK293 cells co-expressing FLAG-D2R, GRK2 and βarrestin2-mCherry enriched in either ethanol (EtOH), DHA or DPA, and incubated with anti-FLAG M1 antibody and stimulated without QPL (- QPL) or with QPL for 10 (+ QPL 10') or 90 minutes (+ QPL 90'). **B**, Quantification of the number of intracellular FLAG-D2R clusters per cell in all 9 conditions from cells as in A. n = 6-22 cells/condition collected across 2 independent experiments. One-way ANOVA followed by Dunnett's multiple comparison test: *** $p<0.001$, ** $p<0.01$, * $p<0.05$, ns $p \geq 0.05$. **C**, Quantification of βarrestin2-mCherry fluorescence in D2R clusters relative to βarrestin2-mCherry fluorescence outside D2R clusters, after stimulation with QPL for 10 (+ QPL 10') or 90 minutes (+ QPL 90'), from cells as in A. n= 6-22 cells/condition collected across 2 independent experiments. Kruskal-Wallis test followed by Dunn's multiple comparisons test: * $p<0.05$, ns $\geq 0.05$.

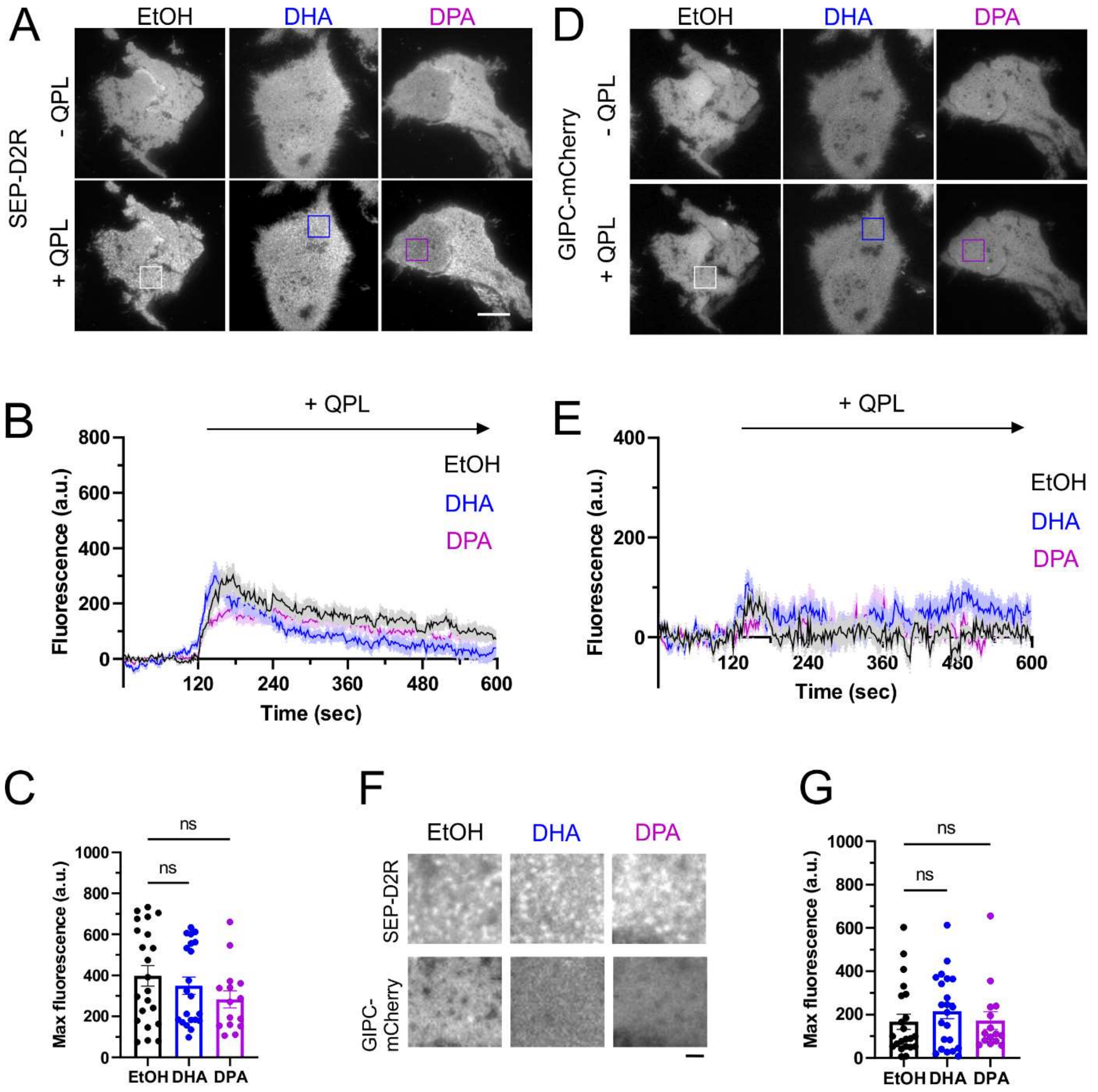


**Figure S6: PUFA enrichment does not affect recruitment of GIPC-mCherry to SEP-D2R clusters. A, D**: Representative TIRF microscopy images of HEK293 cells co-expressing SEP-D2R, GIPC-mCherry, βarrestin2 and GRK2, enriched in either ethanol (EtOH), DHA or DPA, showing the pattern of expression at the plasma membrane for SEP-D2R (A) and GIPC-mCherry (D) before and after stimulation with 10 μM QPL. Scale bar 5 μm. **B, E:** Mean Fluorescence intensity profiles of SEP-D2R (B) and GIPC-mCherry (E) obtained before and after QPL addition at t = 120 s. Data represent average fluorescence values within an ROI drawn around each cluster minus background fluorescence (fluorescence measured in the ROI during the 60 frames before agonist addition was averaged and subtracted from fluorescence values at each frame). **C, G:** Maximum fluorescence intensities of SEP-D2R (C) and GIPC-mCherry (G) measured after QPL addition. Values represent maximum intensity fluorescence minus background fluorescence in each ROI for n= 22, 21 and 15 cells per condition collected across 3 independent experiments. Kruskal-Wallis test followed by Dunn's multiple comparisons test: ns $\geq$0.05. **F:** Zoom-in images taken from panels in A and D, as shown by ROIs, depicting SEP-D2R and GIPC-mCherry clusters after QPL addition. Scale bar 1 μm.

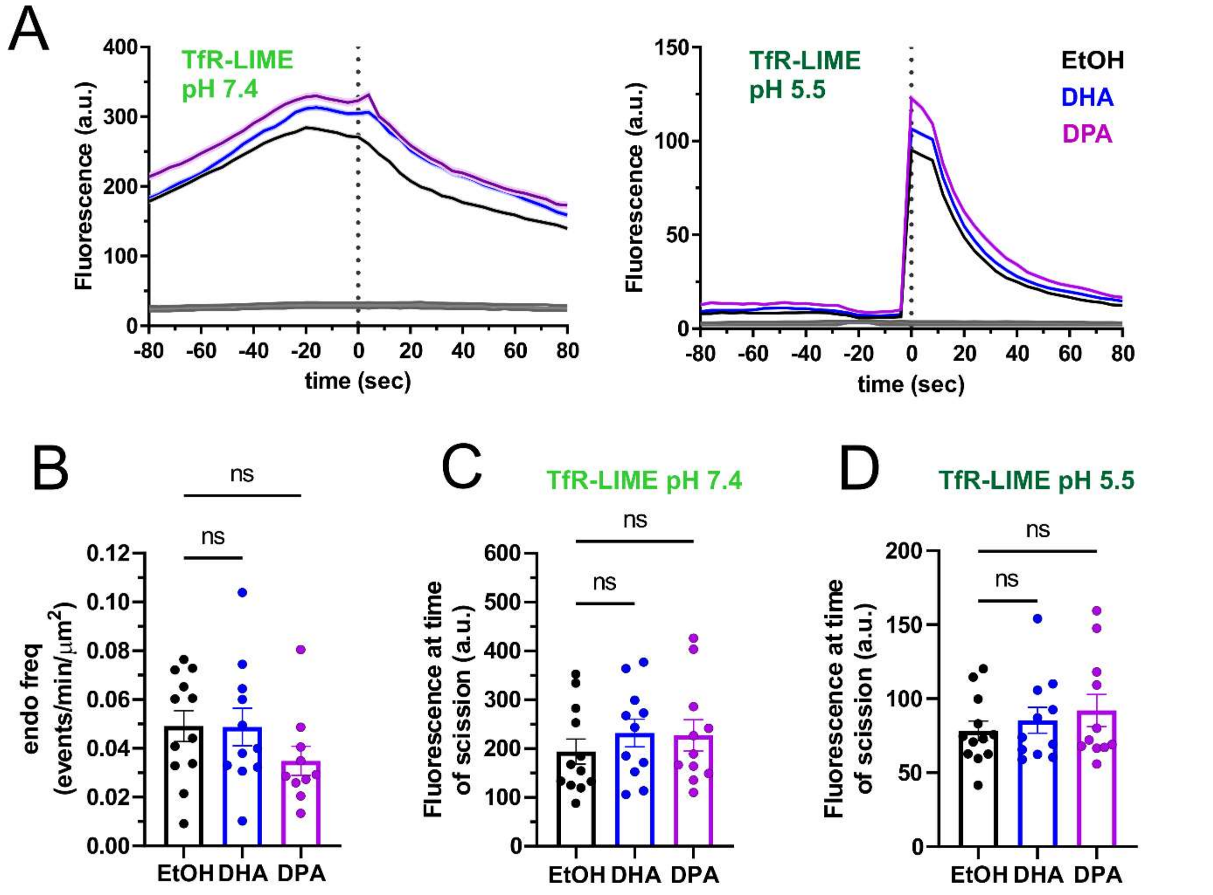


**Figure S7: PUFA enrichment does not affect TfR-SEP endocytosis measured with the ppH assay. A:** Average fluorescence over time of TfR-Lime at pH 7.4 and TfR-LIME at pH 5.5 aligned to the time of vesicle scission (t = 0 s) obtained from terminal events (events for which the TfR-Lime cluster at pH 7.4 disappears within 80 s, defined as in (Taylor *et al*, 2011; Sposini *et al*, 2020) from HEK293 cells transfected with TfR-Lime, treated with either EtOH, DHA or DPA and imaged live with the ppH protocol. **B:** Number of endocytic events/min/$\mu m^2$ detected in the same cells as in A. One-way ANOVA with Dunnett's multiple comparisons test of 12, 11 and 10 cells treated with EtOH, DHA or DPA, respectively; ns $p \leq 0.05$. **C-D:** Average fluorescence intensity of all events in a given cell, at the time of endocytic event detection, for TfR-Lime at pH 7.4 (C) and TfR-Lime at pH 5.5 obtained from the same cells as in A. One-way ANOVA with Dunnett's multiple comparisons test of 12, 11 and 10 cells treated with EtOH, DHA or DPA, respectively; ns $p \leq 0.05$.

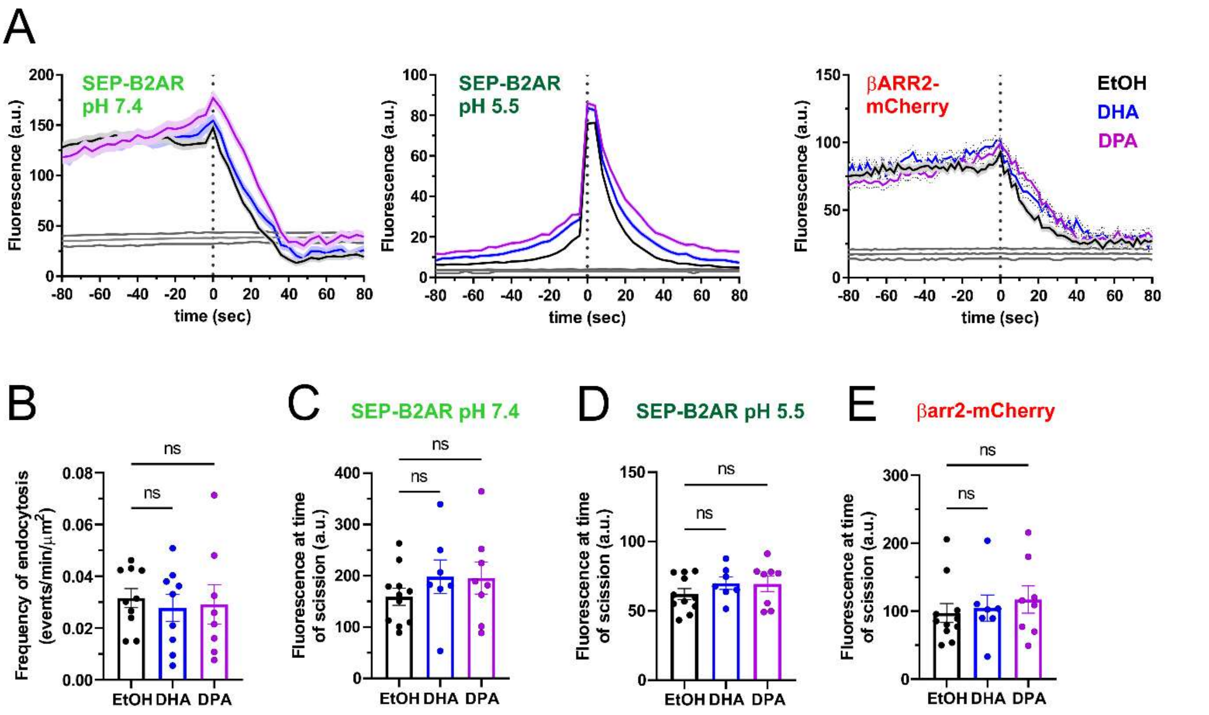


**Figure S8: PUFA enrichment does not affect SEP-β2AR endocytosis induced by Iso and measured with the ppH assay. A:** Average fluorescence over time of SEP-B2AR at pH 7.4, SEP-B2AR at pH 5.5 and β-arrestin2-mCherry at pH 5.5, aligned to the time of vesicle scission (t = 0 s) obtained from terminal events (events for which the SEP-B2AR cluster at pH 7.4 disappears within 80 s, defined as in (Taylor *et al*, 2011; Sposini *et al*, 2020) from HEK293 cells transfected with SEP-B2AR, βarr2-mCherry and GRK2, treated with either EtOH, DHA or DPA and imaged live with the ppH protocol before (0-120 s), during (121-720 s) and after (701-900 s) application of 100 nM Isoproterenol. **B:** Number of endocytic events/min/$\mu m^2$ detected during Isoproterenol application in the same cells as in A. One-way ANOVA with Dunnett's multiple comparisons test of 10, 9 and 8 cells treated with EtOH, DHA or DPA, respectively; ns $p \leq 0.05$. **C-E:** Average fluorescence intensity of all events in a given cell, at the time of endocytic event detection, for SEP-D2R at pH7.4 (C), SEP-D2R at pH 5.5 (D) and β-arrestin2-mCherry at pH 5.5 (D) obtained from the same cells as in A. One-way ANOVA with Dunnett's multiple comparisons test of 10, 9 and 8 cells treated with EtOH, DHA or DPA, respectively; ns $p \leq 0.05$.

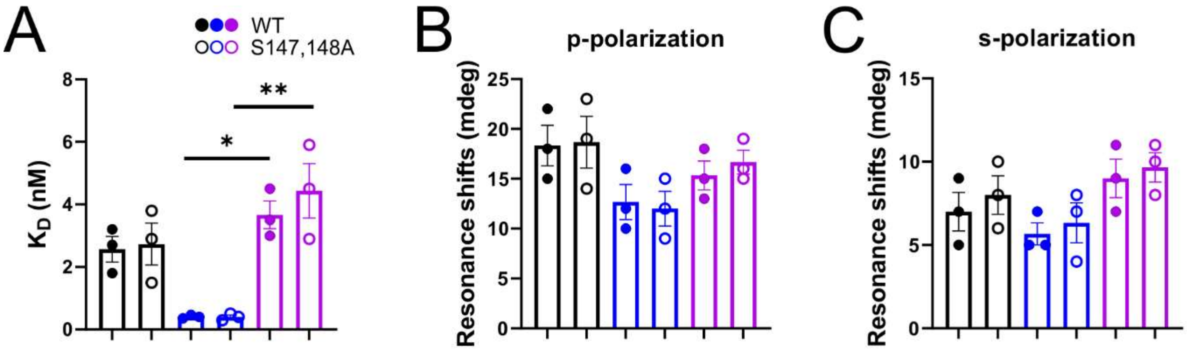


**Figure S9: The modulation of D2R affinity for quinpirole by DHA and DPA is the same for WT and S147, 148A mutant receptors.** **A**, QPL binding affinity to D2R WT (filled circles) and S147,148A mutant in membranes from cells treated with EtOH (black), DHA (blue) or DPA (magenta). One-way ANOVA with Dunnett's multiple comparisons test of WT DHA vs DPA $p = 0.0107$; or mut DHA vs DPA $p = 0.0018$. **B-C**, conformational changes induced by QPL in PUFA-enriched cell membranes observed by PWR with p-polarization (B) and s-polarization (C). Membranes from non-treated cells are presented in black, DHA- and DPA-enriched in blue and magenta, respectively.

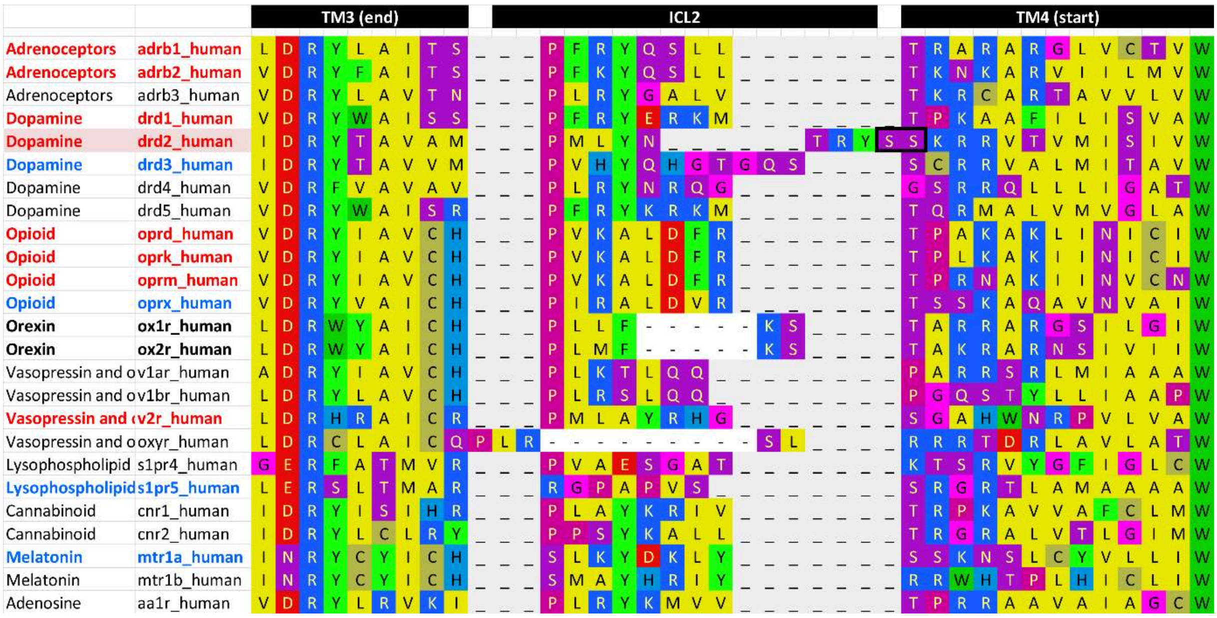


**Figure S10: Alignment of residues in regions TM3, ICL2 and TM4 of a selection of class A GPCRs.** Alignment performed with GPCRdb. Left, agonists and names of human GPCRs (UniProt). In red, the ones tested in this study for their dependency to DHA and DPA enrichment. In blue, receptors containing consecutive serines at the junction between ICL2 and TM4. Black rectangle highlights consecutive serine residues (S) at the end of ICL2 of D2R (S147 and S148). Residues (in one letter code) are in colors corresponding to their chemical properties.

**Movies S1, 2, 3:** Recruitment and clustering of SEP-D2R (left) and β-Arrestin2-mCherry (right) before and during application of QPL observed with TIRF microscopy in cells treated with Ethanol carrier (EV1), DPA (2) and DHA (3)

**Movie S4:** Formation of endocytic vesicles detected with the ppH assay before, during and after application of QPL. SEP-D2R at extracellular pH 7.4 (left) and pH 5.5 (middle), β-Arrestin2-mCherry at pH 5.5 (right).

**Movie S5:** Formation of endocytic vesicles detected with the ppH assay. TfR-Lime at extracellular pH 7.4 (left) and pH 5.5 (right).

**Movie S6:** Formation of endocytic vesicles detected with the ppH assay, before, during and after application of Isoproterenol (ISO). SEP-B2AR at extracellular pH 7.4 (left) and pH 5.5 (middle), β-Arrestin2-mCherry at pH 5.5 (right).
